# DTCFinder: a bench-to-bits toolkit for label-free, whole-spectrum analysis of disseminated and circulating tumor cells in liquid biopsies

**DOI:** 10.1101/2025.11.27.690914

**Authors:** Xiaowei Yan, Zhuo Wang, Ziming Li, Yichun Zhao, Shun Lu, Qihui Shi, Wei Wei

## Abstract

Disseminated (DTCs) and circulating tumor cells (CTCs) are rare but pivotal in understanding cancer metastasis and advancing liquid biopsy-based diagnostics. Traditional detection approaches, hindered by their dependency on epithelial markers or morphological features, impair the accurate identification of varied DTC/CTC phenotypes within liquid biopsies. We describe DTCFinder, a bench-to-bits, label-free method integrating high-throughput single-cell RNA sequencing and machine learning, enabling unbiased detection, in-depth molecular characterization, and precise tissue-of-origin tracing of DTCs/CTCs across various types of liquid biopsies, even with ultra-scarce DTC/CTC presence in peripheral blood and low sequencing depth. DTCFinder offers a robust and versatile toolkit for liquid biopsy-based cancer diagnostics, enhancing our understanding of cancer metastasis.

---

Tumor-derived materials in liquid biopsies, such as disseminated and circulating tumor cells (DTCs/CTCs), circulating tumor DNAs (ctDNAs), and tumor-derived exosomes (TEXs), are garnering increasing recognition for their transformative potential in cancer diagnostics, therapeutic monitoring, and prognosis prediction^1^. CTCs and DTCs, as metastatic precursor cells, detach from primary tumors and disseminate in blood and other body fluids, respectively. Unlike ctDNAs and TEXs, these intact tumor cells not only encapsulate the entire spectrum of molecular information but also have the capability to provide crucial phenotypic and functional insights^2^. This positions DTCs/CTCs as valuable surrogates for primary tumor lesions and as crucial windows into understanding tumor metastasis^3, 4^. Yet, the detection of DTCs/CTCs is fraught with challenges. Current methods, typically reliant on epithelial markers like epithelial cell adhesion molecule (EpCAM) and cytokeratin (CK), or morphological characteristics, suffer from low sensitivity and high false-positive rates due to the phenotypic diversity of DTCs/CTCs and the presence of non-tumor cells with epithelial traits in liquid biopsies^5–8^. Furthermore, traditional approaches fall short in tracing the Tissue-of-Origin and Tumor Types (TOTT) and obtaining accurate, unbiased single-cell transcriptome data of DTCs/CTCs in a high-throughput fashion, crucial for liquid biopsy-based cancer diagnostics and in-depth molecular investigations of these metastatic cells. The research and clinical applications of DTCs/CTCs have long been impeded by their extreme rarity and the lack of an accurate, generic method for their unbiased detection and analysis across different cancer types through a shared oncogenic trait. Such limitations hinder the full exploitation of the vast information potential inherent in DTCs/CTCs, arguably the most informative tumor-derived material obtainable in liquid biopsies.

Cancer fundamentally arises from genomic instability. As a salient feature, somatic copy number alternations (CNAs) are nearly ubiquitous in solid tumors but occur sporadically in normal cells^9, 10^, positioning it as a robust, generic marker for identifying rare DTCs/CTCs in complex normal cell populations. Nevertheless, the extreme rarity of DTCs/CTCs precludes applying single-cell sequencing directly to these samples for genomic elucidation. Emerging computational endeavors leverage machine-learning algorithms to analyze single-cell RNA sequencing (scRNA-seq) data for DTCs/CTCs detection and TOTT tracing^11, 12^. However, these methods have mostly been validated on synthetic datasets, usually abundant in DTCs/CTCs (>100), derived from prior studies and remain untested against real clinical specimens, which are characteristically scarce in DTCs/CTCs. Moreover, some of these algorithms necessitate prior training on specific cancer types^11^, thus circumscribing their applicability in heterogeneous clinical landscapes. Herein, we introduce the Disseminated/circulating Tumor Cell Finder (DTCFinder), a bench-to-bits, label-free method leveraging genome-wide CNAs for unbiased detection, comprehensive molecular profiling, and TOTT tracing of DTCs/CTCs in diverse liquid biopsy specimens across various cancer types.

DTCFinder includes an experimental component to enrich DTCs/CTCs for high-throughput scRNA-seq and a computational pipeline that harnesses an amalgam of machine-learning models and statistical techniques for DTC/CTC identification and analysis (Fig. 1a). Specifically, we employed tetrameric antibody complexes to co-pellet immune cells and platelets along with red blood cells (RBCs) upon density gradient centrifugation, enriching DTCs/CTCs with leftover immune cells amenable for scRNA-seq. Genome-wide CNAs often manifest at early neoplastic stages and act as a pivotal catalyst in both the onset and advancement of malignancies^13^. Analysis of TCGA data revealed that, even at clinical stage I, CNA burdens of tumor cells markedly surpass those of normal cells adjacent to tumors or in circulation, validating CNAs as reliable DTC/CTC markers (Fig. 1b and Extended Data Fig. 1). Utilizing the single-cell transcriptomic data, DTCFinder segments the genome into contiguous genomic bins and applies a Gaussian mixture model (GMM) to infer relative copy numbers for each bin in every single cell. The inference is predicated on the distribution of average gene expression levels of each genomic segment across different cells (see Methods). By establishing the inferred CNAs of a subset of immune cells in the liquid biopsy as the baseline, an outlier detection algorithm identifies bins with anomalous copy numbers and calculates the accumulated number of such outlier bins (AOBs) for each cell. Cells exhibiting significantly elevated AOB values, indicative of extensive genome-wide CNAs, are identified as DTCs/CTCs.

**Fig. 1.**
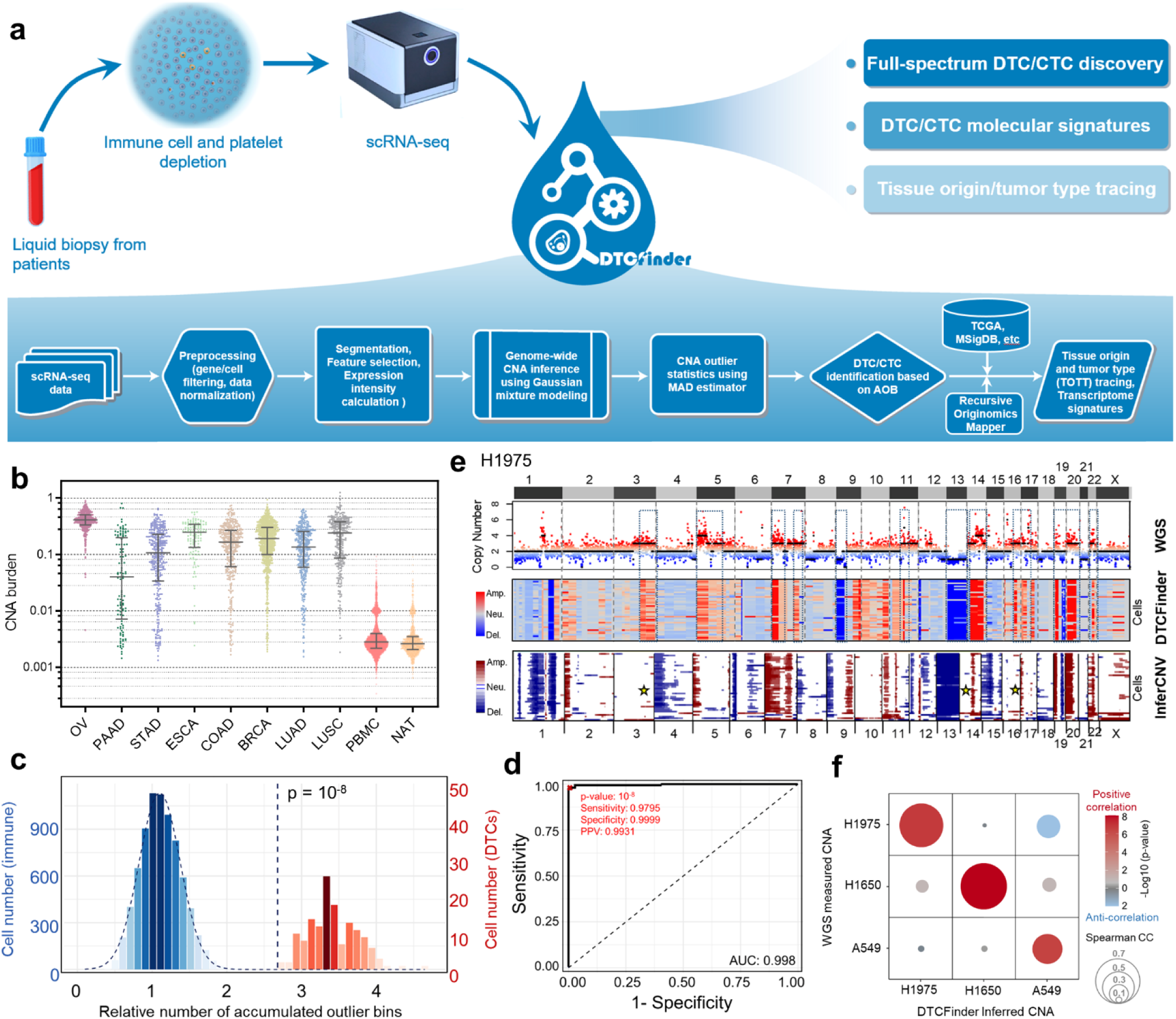
Overview of DTCFinder and core data modalities. **a.** Schematic representation of the integrated experimental and computational workflows encompassed by DTCFinder, with major computational steps highlighted in the flowchart. **b.** Contrasting CNA burdens between normal cells or tissues and eight representative tumor types from TCGA. NAT: Normal Adjacent to Tumor. **c.** Histogram showcasing accumulated outlier bins (AOBs) for baseline immune cells (blue) alongside a Gaussian fitting and spike-in lung cancer cells (red) as discerned by DTCFinder. The dashed line demarcates the AOB cut-off p value of 10^-8^. **d.** ROC curve with sensitivity, specificity, precision (PPV), and AUC values for DTC/CTC identification in the cell line spike-in sample, utilizing the cut-off p value of 10^-8^ (denoted by the red cross in the ROC curve). **e.** Comparative analysis of genome-wide CNAs of spike-in H1975 cells as assessed by LC-WGS against those inferred by DTCFinder and InferCNV. Genomic segments manifesting consistent CNAs are encircled by blue dashed boxes. CNAs at specific genomic loci, accurately inferred by DTCFinder but overlooked by InferCNV, are marked by yellow stars. **f.** Spearman correlations between CNAs measured by WGS and those inferred by DTCFinder across three spike-in lines are represented. The correlation coefficients are symbolized by the circle sizes, and p-values are conveyed through color coding.

The AOB values in immune cells typically follow a Gaussian distribution, and the threshold for DTC identification can be determined from the extreme right tail of this distribution, corresponding to a statistically significant low p-value (Fig. 1c). To empirically ascertain the optimal cut-off p-value, a mixture of three distinct lung cancer cell lines—H1975, H1650, and A549—was spiked into a blood sample, emulating the heterogeneous DTC/CTC populations typically present in liquid biopsy samples, and subsequently subjected to DTC enrichment for scRNA-seq and DTC/CTC identification (Fig. 1c and Extended Data Fig. 2a-d). Utilizing a selective panel of cell line-specific markers, we quantified the cell number of each line in the resulting scRNA-seq dataset (Extended Data Fig. 2e, See Methods). This quantification served as a ground truth for evaluating DTCFinder’s discriminative efficacy. DTCFinder reached highest balance accuracy of 98.98% for the identification of spike-in tumor cells at the cut-off p value of 10^-8^, accompanied by an area under the curve (AUC) of 0.998 and a precision of 99.31% (Fig. 1d). Under p=10^-8^, 150 genes per bin was determined to produce a near-perfect balanced accuracy and the highest precision for DTC/CTC identification (Extended Data Fig. 2f).

To rigorously validate the accuracy of DTCFinder’s inferred genome-wide CNAs, we employed low-coverage whole-genome sequencing (LC-WGS) as a benchmark. Remarkably, the CNAs inferred by DTCFinder were in high concordance with those experimentally measured through LC-WGS across all three lines (Fig. 1e and Extended Data Fig. 2g,h). Furthermore, we conducted a comparative analysis with InferCNV^14^, an existing algorithm for CNA inference. DTCFinder exhibited consistent results with InferCNV, with enhanced precision in certain genomic regions (Fig. 1e and Extended Data Fig. 2g,h). Importantly, unlike DTCFinder, InferCNV requires pre-specified reference cells for CNA inference and lacks the capability for autonomous DTC detection or TOTT tracing.

To ascertain whether DTCFinder’s analytical robustness would be compromised by different types of non-malignant cells, we conducted tests using normal lung tissues and various liquid biopsy samples, including blood and cerebrospinal fluid (CSF) from non-cancer patients^15^. Notably, among these samples were those from patients with multiple sclerosis (MS) containing inflammatory immune cell types. Our findings revealed that normal epithelial cells in lung tissues exhibited AOB distributions that overlapped with CD45+ cells, resulting in no false-positive identifications (Extended Data Fig. 3a,b and Fig. 4). Consistently, no DTC/CTC was identified except for one CSF sample from an MS patient, where 2 false-positive DTCs were found among ∼8000 cells, indicating the high specificity of DTCFinder, even in analyzing complex liquid biopsy samples replete with atypical immune cell types (Extended Data Fig. 3c).

We further extended our analysis by applying DTCFinder to a cohort of liquid biopsy samples obtained from 9 lung cancer patients, 4 of whom had paired primary tumor tissues (Extended Data Fig. 4). The algorithm discerned varying numbers of DTCs/CTCs across different patients. Importantly, the inferred CNAs of DTCs/CTCs closely matched those of paired tumor tissues measured by LC-WGS (Fig. 2a and Extended Data Fig. 5a,b). DTC-specific CNAs in some genomic regions were also observed. Moreover, the inferred CNAs showed marked pairwise correlations across individual DTCs/CTCs, a feature validated by LC-WGS using DTCs/CTCs collected from the same patients (Fig. 2b and Extended Data Fig. 5c,d). Interestingly, in certain samples, such as DTC-S1 and DTC-S2, we identified distinct subgroups of DTCs/CTCs with unique CNA profiles that were highly correlated within the subgroup but less so with other DTCs/CTCs, suggesting the existence of multiple DTC/CTC clones possibly originating from different clonal regions of the primary tumor (Fig. 2b and Extended Data Fig. 5c).

**Fig. 2.**
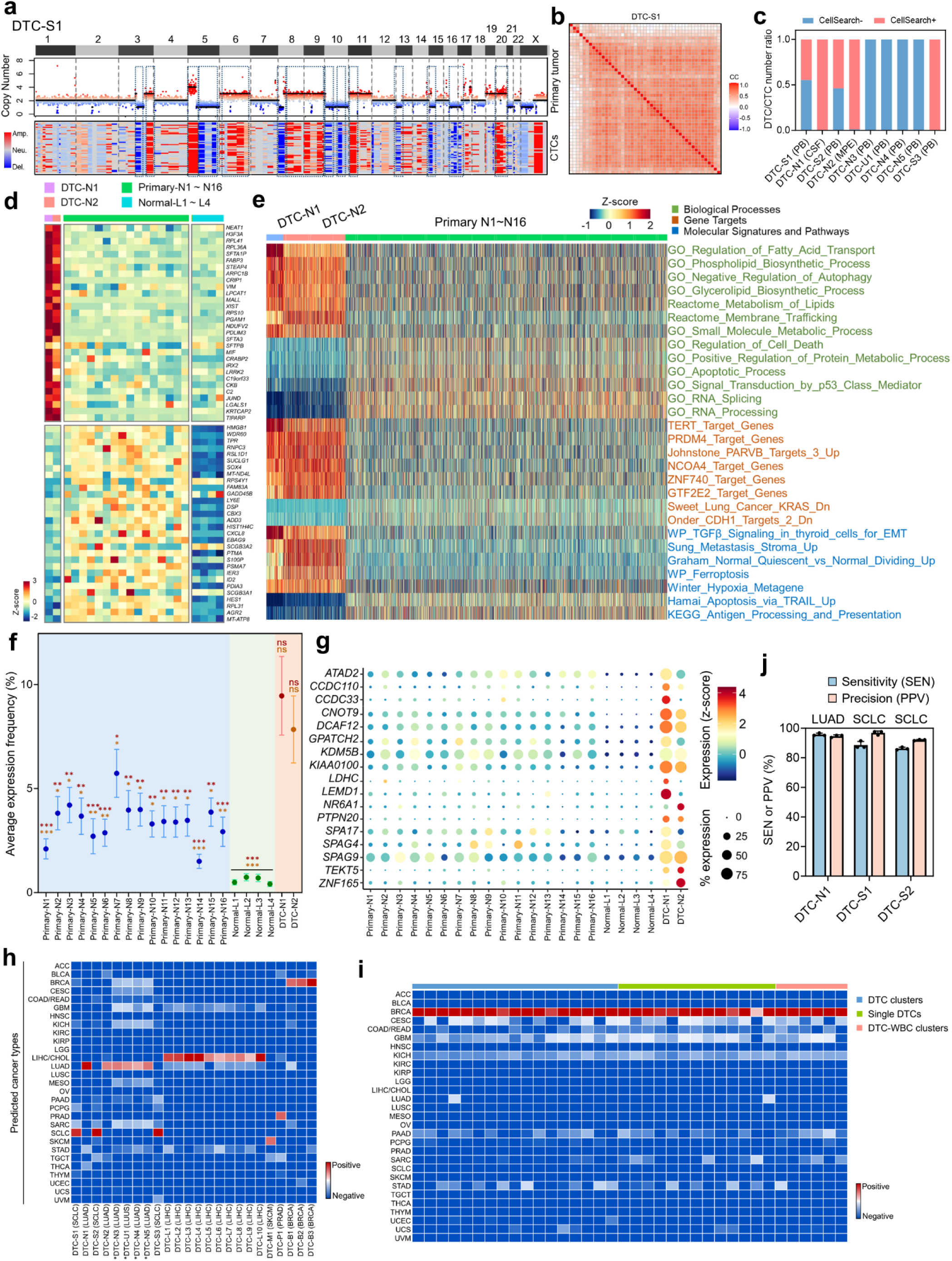
Exemplary analyses of clinical liquid biopsy samples by DTCFinder. **a.** Alignment of genome-wide CNAs of CTCs inferred by DTCFinder with those of the paired primary tumor tissue experimentally measured by WGS for patient DTC-S1. Genomic segments with consistent CNAs are outlined by blue dashed boxes. **b.** A heatmap displaying pairwise Spearman’s correlations of inferred CNAs across all identified DTCs/CTCs in patient DTC-S1 (CC: correlation coefficient). **c.** Proportions of DTCs/CTCs identifiable (CellSearch+) or unidentifiable (CellSearch-) by markers (CD45-/EPCAM+/PanCK+) used in the CellSearch system, across the nine lung cancer patients studied (PB: peripheral blood, MPE: malignant pleural effusion, CSF: cerebrospinal fluid). **d.** Top and bottom 30 genes showing differential expression in DTCs/CTCs from LUAD patients DTC-N1 and DTC-N2, contrasted with primary tumor cells from 16 LUAD patients and 4 normal lung samples. Averaged gene expression levels of each sample are displayed based on the normalized z-scores. **e.** Representative differentially enriched transcriptome signatures between DTCs/CTCs from LUAD patients DTC-N1 and DTC-N2 and primary tumor cells from 16 LUAD patients are color-coded and illustrated with normalized z-scores. For illustrative clarity, the data bars for DTCs/CTCs in the heatmap are enlarged five-fold. **f, g.** Comparative analysis of average expression frequencies across all the cancer-testis antigens (CTA) curated in CTDatabase (**f**), alongside expression levels and frequencies of representative CTAs in DTCs/CTCs from two NSCLC patients (DTC-N1 and DTC-N2), primary tumor cells from 16 LUAD patients, and epithelial cells from 4 normal lung tissues (**g**). In (**f**), mean ± SEM is represented by dots and error bars, with statistical significance assessed via Brown-Forsythe and Welch ANOVA tests, indicated by red (DTC-N1) and orange (DTC-N2) stars for pairwise comparisons, corrected for multiple comparisons using the Benjamini, Krieger, and Yekutieli method (*p<0.05, **p<0.005, ***p<0.0005, ns: not significant). In (**g**), normalized z-scores show averaged CTA expression across single cells within each sample, with circle sizes indicating frequencies (i.e. the percentage of cells expressing the CTA). **h**. Predictions of the tissue-of-origin and tumor type (TOTT) for DTCs/CTCs from patients with different tumor types are color coded as red for positive prediction and blue for negative prediction. The tumor types listed in parentheses next to the sample IDs indicate the diagnosed tumor types. Stars on the sample IDs highlight the samples with 3 or fewer DTCs/CTCs identified. **i.** Prediction of the TOTT for different forms of DTCs/CTCs (color coded by blue, green, and red respectively) identified from a breast cancer patient in a prior study. **j.** The sensitivity and precision of DTC identification using downsampled, low-depth scRNA-seq data from 3 lung cancer patients are depicted across 3 independent downsampling trials (mean ± SD), with accompanying TOTT predictions shown above.

In contrast to FDA-cleared CellSearch system or other similar methods, DTCFinder enables the detection of DTCs/CTCs devoid of epithelial markers. In six out of nine lung cancer patients in our cohort, DTCFinder detected a significant number of EPCAM-negative or PanCK-negative DTCs/CTCs in peripheral blood, undetectable by CellSearch, corroborating previous observations of PanCK-negative DTCs/CTCs in lung cancer patients (Fig. 2c)^8^. Furthermore, given the intrinsic heterogeneity and extreme rarity of DTCs/CTCs, coupled with the presence of confounding cell types in liquid biopsy samples, conventional unsupervised clustering methods prove inadequate for precise DTC/CTC segregation. DTCs/CTCs often do not form an isolated cluster but rather intermingle with confounding cell types, dispersing across multiple clusters within liquid biopsy samples (Extended Data Fig. 6a,b). DTCFinder, capitalizing on hallmark genome-wide CNAs, effectively distinguishes DTCs/CTCs from non-malignant cells, even when they are interspersed within the same clusters. Indeed, for the peripheral blood samples from patient DTC-S1 and DTC-S2, non-CTC cells residing in CTC-enriched clusters were absent from characteristic CNAs present in the CTCs and paired tumor tissues, affirming their non-tumoral nature (Extended Data Fig. 6c).

DTCFinder enables not just the identification but also the comprehensive characterization of DTCs/CTCs by extracting and analyzing their single-cell transcriptome profiles. Through comparison with their matched primary tumors, it delineates DTC/CTC-specific transcriptome signatures, including differentially expressed genes and enriched biological processes and pathways, crucial for unraveling the mechanisms underlying tumor metastasis and identifying targetable vulnerabilities for these metastatic cells. (Fig. 2d,e and Extended Data Fig. 7a-e). For instance, in NSCLC patients DTC-N1 and DTC-N2, we observed elevated expression levels in genes encoding components of the large ribosomal subunit, such as *RPL41* and *RPL36A*, as well as genes associated with a mesenchymal phenotype, like *VIM*, compared with tumor cells from primary lesions or normal lung tissues (Fig. 2d, See Methods). Overexpression of these genes in DTCs/CTCs has been implicated in promoting tumor invasion and metastasis^8, 16^. Consistently, DTCs/CTCs from these patients exhibited elevated signature scores in metastasis and TGFβ signaling pathways while showing reduced scores in apoptosis and antigen-presentation processes, thereby implicating their role in both tumor metastasis and immune evasion (Fig. 2e). Increased metastasis and invasion-related transcriptome signatures were also found in DTCs/CTCs from SCLC patients DTC-S1 and DTC-S2 compared with corresponding primary tumor cells (Extended Data Fig. 7d, e). Interestingly, we detected a marked elevation in both the expression levels and frequencies of many cancer-testis antigens (CTAs) in DTCs/CTCs compared to primary tumor cells or normal lung tissues, indicative of a more aggressive or metastatic phenotype (Fig. 2f,g and Extended Data Fig. 8a,b). Notably, KDM5B, a testis-selective CTA, exhibited increased expression levels and frequencies in DTCs/CTCs from patients with NSCLC and SCLC (Fig. 2g and Extended Data Fig. 8b). The upregulation of KDM5B has been linked to the development of stem-like invasive phenotypes and increased drug tolerance in a variety of cancer types, prompting the active development of pharmacological inhibitors^17^. Some of these CTAs are located in genomic segments with DTC/CTC-specific copy number amplification, which may contribute to the increased expression of these CTAs in the DTCs/CTCs (Extended Data Fig. 8c,d).

Tracing the TOTT is pivotal for leveraging DTCs/CTCs for diagnostic applications, notably in early disease stages where the primary tumor lesion are undetectable via conventional medical imaging or in the monitoring of tumor recurrence. DTCFinder employs its built-in Recursive Originomics Mapper (ROM) module for TOTT tracing using DTC/CTC transcriptome data (Extended Data Fig. 9a). The ROM module commences by formulating tumor type-specific gene signatures from TCGA transcriptome data of 31 solid tumors and develops individual tumor type predictors via logistic regression, optimizing classification accuracy through 10-fold cross validation (Extended Data Fig. 9b, see Methods). Notably, when multiple positive predictions arise for a given sample, we introduce a recursive process to the model to progressively refine the subset until a singular positive TOTT prediction is achieved. This recursive process markedly improved the sensitivity and precision of TOTT predictions across all cancer types (Extended Data Fig. 9c). By applying this prediction module to the 9 lung cancer samples collected in this study, along with 15 additional DTC/CTC samples from published data, DTCFinder demonstrated impeccable prediction accuracy even for samples containing merely a couple of DTCs/CTCs (Fig. 2h and Extended Data Fig. 4). The accurate TOTT tracing even in cases with only a couple of DTCs/CTCs confirms DTCFinder’s capability to precisely pinpoint DTCs/CTCs within samples characterized by extreme DTC/CTC scarcity. We further challenged DTCFinder’s predictive capacity on single CTCs, CTC clusters, and CTC-white blood cell (WBC) clusters where the presence of WBCs may confound the resultant transcriptome data. Using data from a recently published study that collected all three forms of CTC samples from breast cancer patients^18^, DTCFinder rendered 100% accurate predictions across all the samples, confirming the robust TOTT tracing performance of our algorithm (Fig. 2i).

We further assess DTCFinder’s proficiency in contexts with low sequencing depths. We down-sampled the raw scRNA-seq data of three patients, characterized by relatively high number of DTCs/CTCs, from 34,000 – 116,000 reads/cell to 5000 reads/cell to quantitatively evaluate the influence of sequencing depth on the sensitivity and precision of DTC/CTC identification (See Methods). Remarkably, even under conditions of markedly reduced sequencing depth, DTCFinder maintained its accuracy in DTC/CTC identification and relatively consistent inferred genome-wide CNA profiles, achieving an average precision of 94.53±2.24% and sensitivity of 90.20±4.47% across the three patients (Fig. 2j and Extended Data Fig. 10a). Notably, the identified DTCs/CTCs, even with low-depth transcriptome data, could be precisely traced back to their respective TOTT, underscoring the robust performance of DTCFinder on low-depth scRNA-seq data (Extended Data Fig. 10b).

In summary, DTCFinder offers a bench-to-bits toolkit for label-free, full-spectrum DTC/CTC discovery and analysis across various cancer types. It encompasses a holistic view of DTCs/CTCs, ranging from genome-wide CNAs and transcriptome signatures to TOTT tracing, demonstrating versatility across diverse types of liquid biopsy and compatibility with multiple commercial scRNA-seq platforms without a demand for deep sequencing, The adaptability of DTCFinder, positions it as a useful tool, readily adoptable by research community for fundamental and clinical studies on liquid biopsy-based cancer diagnosis and cancer metastasis.

## Methods

### Patient information and sample collection

Liquid biopsy samples including peripheral blood, malignant pleural effusion (MPE), and cerebrospinal fluid (CSF) in this study were obtained from Shanghai Chest Hospital between November 2019 and July 2022. The protocols were performed according to the principles of the Helsinki Declaration and approved by the Institutional Review Board (#KS1973, #IS21109).

### Cell lines and reagents

Lung cancer cell lines H1975, H1650 and A549 were obtained from American Type Culture Collection (ATCC). Cell lines were routinely maintained in ATCC-formulated cell culture medium containing 10% fetal bovine serum (FBS, Gibco) and 1× Penicillin-Streptomycin-Glutamine (Gibco) in a humidified atmosphere of 5% CO_2_ and 95% air at 37°C. Reagents used in this study have been listed in Table 1 below.

**Table 1.**
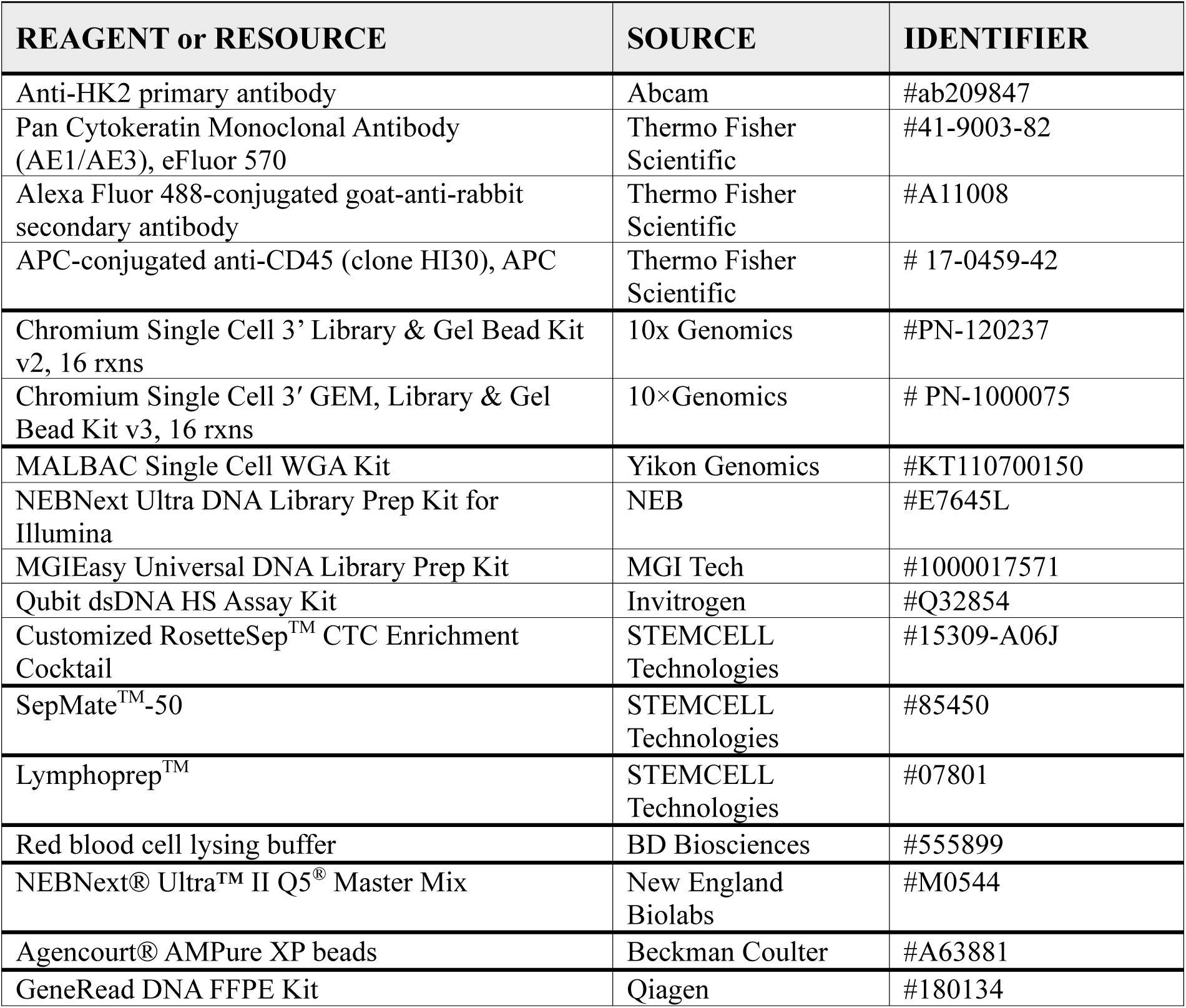
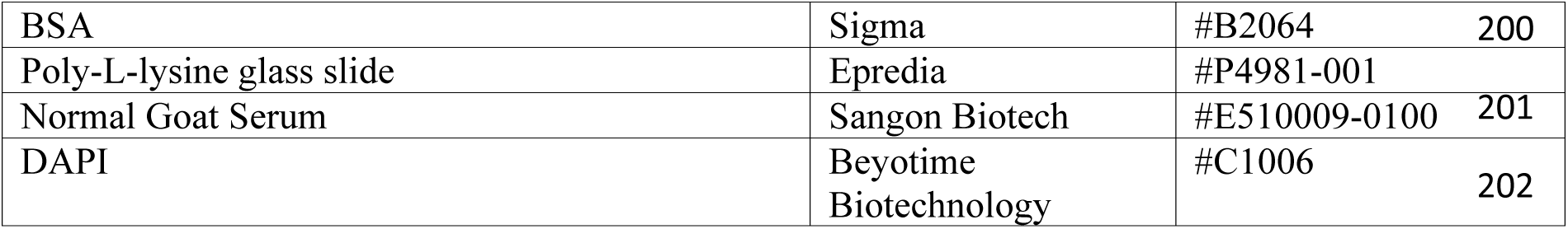
List of reagents.

| **Table 1 \| List of reagents** |  |  |
| --- | --- | --- |
| **REAGENT or RESOURCE** | **SOURCE** | **IDENTIFIER** |
| Anti-HK2 primary antibody | Abcam | #ab209847 |
| Pan Cytokeratin Monoclonal Antibody (AE1/AE3), eFluor 570 | Thermo Fisher Scientific | #41-9003-82 |
| Alexa Fluor 488-conjugated goat-anti-rabbit secondary antibody | Thermo Fisher Scientific | #A11008 |
| APC-conjugated anti-CD45 (clone HI30), APC | Thermo Fisher Scientific | # 17-0459-42 |
| Chromium Single Cell 3' Library & Gel Bead Kit v2, 16 rxns | 10x Genomics | #PN-120237 |
| Chromium Single Cell 3' GEM, Library & Gel Bead Kit v3, 16 rxns | 10×Genomics | # PN-1000075 |
| MALBAC Single Cell WGA Kit | Yikon Genomics | #KT110700150 |
| NEBNext Ultra DNA Library Prep Kit for Illumina | NEB | #E7645L |
| MGIEasy Universal DNA Library Prep Kit | MGI Tech | #1000017571 |
| Qubit dsDNA HS Assay Kit | Invitrogen | #Q32854 |
| Customized RosetteSep™ CTC Enrichment Cocktail | STEMCELL Technologies | #15309-A06J |
| SepMate™-50 | STEMCELL Technologies | #85450 |
| Lymphoprep™ | STEMCELL Technologies | #07801 |
| Red blood cell lysing buffer | BD Biosciences | #555899 |
| NEBNext® Ultra™ II Q5® Master Mix | New England Biolabs | #M0544 |
| Agencourt® AMPure XP beads | Beckman Coulter | #A63881 |
| GeneRead DNA FFPE Kit | Qiagen | #180134 |

|  |  |  |  |
| --- | --- | --- | --- |
| BSA | Sigma | #B2064 | 200 |
| Poly-L-lysine glass slide | Epredia | #P4981-001 |  |
| Normal Goat Serum | Sangon Biotech | #E510009-0100 | 201 |
| DAPI | Beyotime Biotechnology | #C1006 | 202 |

### DTC/CTC enrichment and single-cell RNA sequencing

Peripheral blood (5 ml), MPE (10 ml) and CSF (1 ml) were collected and sent to the lab at 4^°^C immediately after collection for sample processing and high-throughput scRNA-seq. MPE samples were filtered by a membrane with a pore size of 70 μm and centrifuged at 300 g for 10 min to separate cell pellets. Cells were then treated with red blood cell (RBC) lysing buffer (BD Biosciences) to remove RBCs, followed by depletion of leukocytes with EasySep^TM^ Human CD45 Depletion Kit II (STEMCELL Technologies). The cell pellets were re-suspended in 0.5 mL of PBS. CSF samples were centrifuged at 300 g for 10 min to separate cell pellet, and re-suspended in 0.5 mL of PBS without further enrichment. Peripheral blood samples were centrifuged at 500 g for 5 min, and the cell pellets were re-suspended in an equivalent volume of HBSS and mixed with 75 μl tetrameric antibody cocktail (customized RosetteSep^TM^ CTC Enrichment Cocktail, STEMCELL Technologies) at room temperature for 20 min, followed by adding 15 ml of HBSS with 2% FBS and mixing well. The mixture was carefully added along the wall of the Sepmate tube (SepMate^TM^-50) after adding 15 ml density gradient liquid (Lymphoprep^TM^) into the tube through the middle hole. After centrifuging at 1200 g for 20 min, the topmost supernatant (∼10 ml) was discarded, and the remaining liquid (∼10 ml) above the barrier of the Sepmate tube was rapidly poured out into a new centrifuge tube with addition of 20 ml of HBSS. After centrifuging at 600 g for 8 min, the supernatant was removed and 1 ml of RBC lysing buffer was then added for 5 min to lyse RBCs. After centrifuging at 450 g for 5 min, the nucleated cell pellet was re-suspended in 100 μl of HBSS for single-cell RNA sequencing.

High-throughput scRNA-seq was conducted with a target recovery of 8,000 cells. Cell counting and cell viability were measured with a Countess II Automated Cell Counter, and the cell concentration was adjusted to 1000 cells/μL. Cells were loaded into a Chromium Single-Cell 3 Chip Kit v2 and v3 (10× Genomics). Reverse transcription, cDNA recovery, cDNA amplification, and library construction were performed using Chromium Next GEM Single Cell 3’ Reagent Kits v2 and v3 (10x Genomics) according to manufacturer’s instructions. Single-cell library sequencing was performed using the Illumina NovaSeq 6000 platform, with 150 bp paired-end reads.

### Isolation of genomic DNA from primary tumors and copy number alternation profiling

Two serial sections from each individual formalin-fixed and paraffin-embedded (FFPE) tumor tissue block were used to isolate genomic DNA (gDNA) by GeneRead DNA FFPE Kit (Qiagen) according to manufacturer’s instructions. Concentrations of gDNA were quantified with Qubit dsDNA HS Assay Kit (Invitrogen). WGS library was constructed with the NEBNext® Ultra™ DNA Library Prep Kit for Illumina (New England Biolabs) according to manufacturer’s protocols. Libraries were analyzed by Illumina Navoseq 6000 platform with 150 bp pair-end reads (Genewiz, China).

### CTC identification and retrieval, whole genome amplification, and copy number alternation profiling for patient sample DTC-S1

After blood sample processing and CTC enrichment, the enriched cell suspension was applied onto a 3% BSA-treated poly-L-lysine glass slide. Cells on the slide were fixed (2% PFA, 10 min), permeabilized (0.5% Triton X-100, 15 min) and blocked with a solution containing 3% BSA and 10% Normal Goat Serum for 1 hour. On-chip immunostaining was then performed by incubation with APC-conjugated mouse anti-human CD45 antibody, eFluor 570-conjugated mouse anti-human PanCK antibody, and rabbit anti-human hexokinase 2 (HK2) antibody in PBS overnight at 4°C. After extensive washing with PBS, cells on the chip were treated with Alexa Fluor 488-conjugated goat-anti-rabbit secondary antibody in PBS for 1 h and DAPI for 10 min followed by washing with PBS. ImageXpress Micro XLS Wide field High Content Screening System (Molecular Devices) scanned the chip and imaged all cells in bright field and fluorescent channels (CD45: CY5; HK2: FITC, CK: PE, Nucleus: DAPI). A computational algorithm analyzed the images to identify CTCs adhering to previously validated criteria we reported before^8^. Specifically, it identified putative CTCs characterized by distinct marker profiles, including DAPI^pos^/CD45^neg^/HK2^high^/CK^pos^, DAPI^pos^/CD45^neg^/HK2^high^/CK^neg^, and DAPI^pos^/CD45^neg^/HK2^low^/CK^pos^ cells, based on the calculated fluorescence cutoffs generated from HK2 and CK fluorescence intensities of CD45^pos^ leukocytes in the same sample.

Single-cell LC-WGS was used to characterize genome-wide CNA profiles. Single-cell whole genome amplification (WGA) was firstly conducted with the MALBAC® Single Cell WGA Kit (Yikon Genomics). To assess the WGA coverage of amplified product, 22 primer pairs were designed to target 22 loci located on different chromosomes. Six primer pairs were randomly selected for PCR and a successful amplification of at least four out of six primer pairs generated a positive quality control (QC)-PCR. WGA products that passed QC-PCR were then used to construct WGS library with the NEBNext® Ultra™ DNA Library Prep Kit for Illumina (New England Biolabs) or MGIEasy Universal DNA Library Prep Kit (MGI Tech) according to manufacturer’s protocols. The concentrations of purified fragmented DNA or libraries were quantified with Qubit dsDNA HS Assay Kit (Invitrogen). Libraries were sequenced by Illumina Navoseq 6000 with 150 bp pair-end reads (Genewiz, China) or MGI2000 sequencer with 100 bp single-end reads (JunHealth, China).

### Pre-processing of single-cell RNA sequencing data

FASTQ files of patient samples were individually processed to generate a feature-barcode matrix (genes × cells) using CellRanger version 6.1.2. The processing was performed with the refdata-cellranger-GRCh38-3.0.0 human genome reference, employing default parameters tailored for the 10X single-cell 3’ library and setting expected cell counts to 10,000. Genuine cells were distinguished from empty droplets based on the cumulative distribution of total transcript counts. Cells exhibiting low library complexity, characterized by either gene counts less than 200 or total transcript counts less than 500 were filtered out. Additionally, cells displaying a substantial fraction of mitochondrial transcripts, exceeding 1/3 of their content, were excluded. Prior to inferring single-cell CNAs using DTCFinder, erythroid cells were also removed.

### Construction and implementation of the DTCFinder algorithm

The overall workflow of DTCFinder is depicted in Fig. 1a and includes the following steps

#### (1) Data normalization and transformation

Each cell’s feature UMI counts were normalized by dividing by its total UMI counts, scaling by a factor of 10^5^. After this normalization, genes were retained for subsequent analysis if their total normalized counts summed over all cells exceeded 10. Finally, the normalized UMI counts were log2-transformed, with a pseudocount of 1.

#### (2) Segmentation, feature selection, and expression intensity calculation

The entire human genome (GRCh38.p13) was sequentially segmented by chromosomes, ensuring each segment (or bin) exclusively encompassed a pre-defined number of genes. A default of 150 genes per bin was chosen based on its optimal balanced accuracy and minimal FDR (Extended Data Fig. 2f). Gene positional information was procured using the R package biomaRt^19^, referencing the ENSEMBL_MART_ENSEMBL dataset within the BioMart database and the hsapiens_gene_ensembl dataset. For every genomic bin per cell, the expression intensity was computed as the mean expression of the encompassed 150 genes, centralized to the cell’s average gene expression, and subsequently utilized for CNA inference.

#### (3) Genome-wide CNA inference using Gaussian mixture modeling

To discern genome-wide CNAs for individual cells, a GMM was implemented. Given that normal cells and DTCs/CTCs in liquid biopsy samples have distinct CNA signatures, a dual-component GMM was computed for each genomic segment over the calculated expression intensities. Explicitly for each bin, the Expectation-Maximization (EM) algorithm acted on the z-score standardized expression intensities across cells, leveraging the R package mixtools^20^, to determine maximum likelihood estimates and posterior probabilities. The posterior probabilities, as derived from the EM algorithm for Gaussian distribution mixtures, were subsequently utilized to simulate the cells’ CNAs over each genomic bin^21^. This simulation procedure was reiterated eight times, selecting the GMM that maximized output likelihood estimates. Convergence of the EM algorithm was confirmed if the data log-likelihood varied by less than ε = 10^-8^ during iteration and was deemed divergent if iterations exceeded 1000 times. Any genomic bin manifesting consistent divergence across all 8 simulations was excluded from further scrutiny. This methodology culminated in a posterior probability matrix (cells × bins) for single cells over the genomic bins modeled with definitive GMMs. These posterior probabilities were z-score standardized across cells and can be visualized as a heatmap post-standardization using pheatmap 1.0.12 package^22^. Given that in liquid biopsies, immune cells and other normal cells considerably outnumber DTCs/CTCs and manifest homogenous CNA patterns, their z-scored posterior probabilities are approximated to zero, barring occasional outlier gene expressions in specific cells. This represents the diploid copy number baseline. In contrast, DTCs/CTCs typically display pronounced z-scores in altered chromosomal segments due to substantial expression variations of genes in those bins with respect to normal immune cells, highlighting potential amplified or deleted chromosomal regions.

#### (4) Copy number outlier statistics and DTC identification

DTCs/CTCs typically exhibit much more pronounced genome-wide CNAs compared to normal cells. The extent of these CNAs across the genome aids in DTC identification. To establish a baseline copy number profile for each genomic bin, we utilized the posterior probabilities of CD45+ immune cells (excluding those with PTRPC expression levels within the first quartile of all CD45+ cells). The median absolute deviation (MAD) of these posterior probabilities was then computed for each bin. The copy number deviation of each cell from this baseline for a specific genomic bin was determined by calculating the difference between its posterior probability and the sample median of the posterior probabilities for that bin. If this deviation exceeded 5×MAD, the cell was classified as a copy number outlier for that particular bin^23^. Subsequently, we quantified the degree of genome-wide CNAs for each cell by tallying the number of bins identified as outliers. The accumulated outlier bins (AOBs) for the baseline immune cells were minimal and conformed to a Gaussian distribution. Conversely, DTCs/CTCs typically displayed significantly heightened numbers of AOBs. These can be pinpointed from the extreme right tail of the Gaussian distribution, correlating with a notably low p-value indicative of statistical significance. A cell was designated as a DTC if its AOB value surpassed the threshold set by a p-value of 10^-8^ relative to the Gaussian distribution of baseline immune cells (Fig. 1c).

#### (5) Recursive Originomics Mapper for tracing DTCs/CTCs’ tissue-of-origin and tumor type

To ascertain the TOTT of DTCs/CTCs, we devised a machine learning-based Recursive Originomics Mapper (ROM) module, constructing 31 classifiers corresponding to 31 distinct solid tumor types. This model was trained using an integrated transcriptomic dataset, which included 30 solid tumor transcriptomes (comprising 8880 samples) from TCGA and an SCLC dataset from a previous study^24^, inclusive of 81 human SCLC samples across various subtypes.

For each tumor type, we curated a unique gene signature. This signature was formulated by leveraging the integrated transcriptomic data to pinpoint genes manifesting elevated expression in a specific tumor type while maintaining subdued expression in other types. To homogenize these diverse transcriptomic datasets and negate batch effects, we started with Illumina HiSeq percentile datasets sourced from the TCGA hub (UCSC Xena). These datasets rank gene RSEM values within a 0% to 100% spectrum for each specimen. The SCLC datasets were similarly transformed into percentile ranks, mirroring the methodology applied to the TCGA data. Subsequently, a signature gene set was curated for each tumor type (Supplementary Table 1). This was achieved by selecting genes that met the following differential expression criteria in the tumor type of interest: (1) the gene’s average percentile rank across all samples of the target tumor type exceeded 50%, (2) this percentile was at minimum double the average percentile of that gene across samples from other tumor types, and (3) a statistical significance threshold of p < 10^-20^ was met, as determined by the Mann-Whitney U test.

Having derived the gene signature set for each tumor type, we computed a signature score for every sample within the integrated transcriptomic dataset by averaging the sample percentile expressions for all signature genes. Consequently, each sample was endowed with a unique signature score for each of the 31 tumor types, leading to an aggregate of 31 scores for every sample. By extending this computation to the entire sample set, we generated a matrix delineating samples against tumor-type scores. Using this matrix as input, we formulated and validated a classifier for each tumor type. This was executed via logistic regression modeling, employing the R package Epi 2.47^25^ and harnessing the iteratively reweighted least squares (IRLS) methodology for deducing maximum likelihood estimates during the model fitting. To counteract potential overfitting, we instituted a constraint on feature (tumor type) selection, mandating that the number of features in each classifier be capped at five. For every tumor type, its classifier incorporated the signature score of that tumor alongside scores of up to four other tumor types. This integration resulted in a myriad of tumor type combinations for each classifier. For every tumor type’s classifier, model training and parameter tuning were executed through a repeated cross-validation approach. We elected the combination boasting the highest balanced accuracy when distinguishing a specific tumor type from the cumulative 8961 samples spanning the 31 solid tumor categories. These refined 31 classifiers collectively formed the core prediction unit (CPU) of our ROM module (Extended Data Fig. 9a, Supplementary Table 2).

With the CPU in place, the algorithm processes a given sample’s transcriptomic data, determines signature scores for the 31 solid tumor types, and employs the CPU to predict the sample’s TOTT. A sample is designated a particular tumor type (i.e. positive prediction) if its type-specific logit ≥ 0. Owing to the overlapping transcriptomic signatures observed between liver (LIHC) and bile duct (CHOL) cancers, as well as between colon (COAD) and rectal (READ) cancers, our algorithm consolidates these analogous cancer types into a unified class for predictive purposes. If a singular positive prediction appears, that specific cancer type is reported. In the absence of positive predictions, the outcome is labeled “undetermined”. However, for scenarios with multiple positive predictions, the algorithm initiates a recursive process.

In this recursive phase, the ROM module redefines the signature gene sets and logistic regression model solely for the tumor types positively identified in the initial round. Utilizing only the transcriptomic data of these specified tumor types from the integrated transcriptomic dataset, the algorithm recalibrates gene signatures based on our earlier criteria, computes updated signature scores, and then reconstructs the classifiers via logistic regression modeling using these refreshed scores. Using this overhauled CPU, the algorithm then revisits the task of determining the sample’s TOTT. If a singular positive prediction emerges, that specific cancer type is reported. Otherwise, if no new predictions or the same number of positive predictions as the previous round arise, the cancer types identified in the initial round are reported. Should a reduced number of positive predictions manifest, the recursive procedure is invoked once more, refining the signature gene sets and logistic regression models tailored to the tumor types positively predicted in the second round. This iterative process persists until a definitive prediction is secured or until the predictive outcomes stabilize (Extended Data Fig. 9a).

To gauge our ROM module’s efficacy, we undertook a 10-fold cross-validation on the integrated transcriptomic dataset encompassing the 31 solid tumor types. In this evaluation, samples were arbitrarily segmented into ten equally sized groups. The model was trained using nine of these groups and tested on the remaining one, cycling through all groups until each had served as the testing dataset. This validation yielded a sensitivity of 92.9±5.8% and a precision of 94.8±6.4%, attesting to the model’s robust performance (Extended Data Fig. 9b). Our finalized model, trained on the entire transcriptomic dataset of the 31 tumor types, was then primed for predicting DTCs/CTCs’ TOTT. Using our algorithm, DTC/CTC transcriptomic data from a specific sample underwent preprocessing described above, with gene expression levels averaged across all discerned DTCs/CTCs to fashion a percentile-ranked dataset. Leveraging this data, our algorithm determined signature scores for the 31 solid tumor types and employed the established ROM module to predict that sample’s TOTT.

#### (6) Implementation of DTCFinder

DTCFinder, designed for the R software environment, offers straightforward installation and operation through simple command-line interfaces. This algorithm boasts compatibility with a diverse array of data input formats and provides comprehensive outputs, including the identification of DTCs/CTCs, their genome-wide CNA profiles, clonal compositions, transcriptome profiles, and TOTT tracing results. The software package, along with a detailed user guide and demonstrative examples, is available for download upon publication.

### Visualization and clustering of single-cell RNA sequencing data

To test whether unsupervised clustering can directly separate DTCs/CTCs into distinct clusters (Extended Data Fig. 6a,b), we employed Seurat to process, visualize, and cluster scRNA-seq data from patient samples DTC-S1 and DTC-S2^26^. These specific samples were chosen due to their origin from peripheral blood, containing a relatively large number of DTCs/CTCs, which aptly represent the heterogeneity of DTCs/CTCs in circulation and provide a sufficient sample size of DTCs/CTCs for clustering analysis. Unless otherwise specified, default parameters were maintained throughout the process. Raw count data underwent normalization in Seurat using the “LogNormalize” approach followed by the selection of top 10000 variable features using vst method and data scaling and centralization over these selected features. We then used UMAP projections to generate lower dimensional representations on the first 50 principal components of the chosen features using knn = 20. Original Louvain algorithm was used to optimize the modularity function to determine clusters.

### Copy number alteration analysis from low-coverage whole genome sequencing data

FASTQ files were aligned to the major chromosomes of human (GRCh38) using bowtie2-2.3.5.1 with default parameters^27^. PCR duplicates were removed with Samtools (version 1.11)^28^. Aligned reads were counted in fixed bins averaging 500 kb. Bin counts were normalized for GC content with lowess regression and bin-wise ratios were calculated by computing the ratio of bin counts to the sample mean bin count. The diploid regions were determined using HMMcopy (version 0.1.1)^29^. Segmentation was performed with circular binary segmentation (CBS) method (alpha = 0.0001 and undo.prune = 0.05) from R Bioconductor DNAcopy package^30^. Copy number noise was quantitated using the mean absolute pairwise difference (MAPD) algorithm. Samples with MAPD ≤ 0.45 passed the MAPD QC and were included in single-cell genome-wide CNA analyses.

### Identification of spiked-in cancer cells via cell line-specific markers

To accurately discern and quantify spiked-in H1975, H1650, and A549 cells, thereby establishing a benchmark for evaluating DTCFinder’s classification precision, we derived cell line-specific signature genes from existing RNA-seq profiles of non-small cell lung cancer cells^31^. This dataset (GSE160683) features deep RNA sequencing of 60 human lung cell lines alongside a human lung epithelial cell line and two human lung fibroblast cell lines, each with three replicates. Differentially expressed genes (DEGs) between H1975 cells and an ensemble of other cell lines, including H1650, A549, BEAS-2B (lung epithelial cells), IMR-90 (lung fibroblasts), and WI-38 (lung fibroblasts) cells, were discerned. The criteria for DEGs were an expression fold change > 2 and p < 0.05 as determined by a two-tailed Welch’s t-test. Following this, the DEGs were ranked by the magnitude of expression fold change. A manual evaluation of the top 30 ranked DEGs against all other lung cancer cell lines in the dataset was conducted. This led to the selection of 7 signature genes as shown below, characterized by pronounced expression in H1975 cells and minimal expression in H1650, A549, BEAS-2B, IMR-90, WI-38, and other lung cancer cell lines. An additional requirement for these signature genes was negligible expression in immune cells, as verified via The Human Protein Atlas (https://www.proteinatlas.org/). Using a similar methodology, seven signature genes for H1650 and A549 were also identified, respectively. When clustering the expression matrix of the 21 identified signature genes across all single-cell transcriptome data of the spiked-in sample, the spiked-in H1975, H1650, and A549 cells distinctly clustered into three separate groups, clearly demarcated from immune cells (Expanded Data Fig. 2e).

A549 signature genes: *RSPO3, ARG2, INSL4, GABRB3, GABRA5, HOXB8, NF0B1*

H1650 signature genes: *SPINK6, MAL2, LY6D, CBLC, VGLL1, CSAG1, MAGEA12*

H1975 signature genes: *KCNIP3, CCDC3, IFITM10, DHRS2, ZNF560, P2RX6, CT83*

### Analysis of normal lung tissues and liquid biopsy samples from noncancerous patients

To gauge the false positive rate of DTCFinder when evaluating normal tissue or liquid biopsy samples from individuals without cancer, we examined scRNA-seq datasets derived from two prior studies^15, 32^. These datasets encompass scRNA-seq data from four freshly excised human lung tissues, specifically parenchymal lung and distal airway specimens, taken from uninvolved sections during tumor resections in 4 patients. Additionally, the datasets include scRNA-seq data from 22 CSF and PBMC samples harvested from 6 multiple sclerosis (MS) patients and 6 idiopathic intracranial hypertension (IIH) patients. Following the preprocessing steps described above, the data underwent analysis using DTCFinder. In this evaluation, a mere 2 out of approximately 8,000 cells from a CSF sample of an MS patient were incorrectly identified as DTCs (Extended Data Fig. 3c). Notably, the AOBs of these misclassified cells were marginally proximate to the cut-off AOB threshold, defined by a p-value of 10^-8^.

### Analysis of differential transcriptome signatures and cancer-testis antigens between DTCs/CTCs and matched primary tumor cells

To discern differential transcriptomic signatures between DTCs/CTCs and the primary tumor cells from the same cancer types, we sourced scRNA-seq datasets for NSCLC and SCLC from two prior studies^33, 34^. Specifically, scRNA-seq data was extracted from 16 LUAD patient tumors (Primary-N1-N16) and 19 SCLC patient tumors spanning various subtypes (Primary-SA1-SA13 and Primary-SN1-SN6) as detailed in Extended Data Fig. 4. For LUAD samples, UMI count data from LUAD patient samples DTC-N1 and DTC-N2 were amalgamated with data from primary tumor cells of 16 LUAD patients (Primary-N1-N16). In the SCLC dataset, subtype determination for samples DTC-S1 and DTC-S2 was performed based on marker gene expression for *ASCL1*, *NEUROD1*, *POU2F3*, and *YAP1*. DTC-S1 was classified as SCLC-N (*NEUROD1+*), and DTC-S2 as SCLC-A (*ASCL1+*) (Extended Data Fig. 7a). Thereafter, UMI count data for sample DTC-S1 was paired with data from SCLC-N patient tumors (Primary-SN1-SN6), and for sample DTC-S2 with SCLC-A tumors (Primary-SA1-SA13).

The integrated datasets underwent preprocessing as previously described, followed by normalization by library sizes, scaling by a factor of 10,000, and natural log-transformation with a pseudocount of 1. Batch effect was assessed for the combined datasets to ensure normalized expression of immune cells of shared phenotypes well-mixes across samples. Cell type annotations for primary tumor samples were applied to identify tumor cells following the same procedures used in the original studies^33, 34^.

DEGs between DTCs/CTCs and their matched primary tumor cells was ascertained using the Seurat 4.3.0 package^26^, with significance determined by the non-parametric Wilcoxon rank-sum test. The top 30 DEGs for both DTCs/CTCs and primary tumor cells, exhibiting a significance level of p<10^-30^, were visualized via a heatmap alongside epithelial cells from the aforementioned normal lung samples (Normal-L1-L4) using pheatmap 1.0.12 package^22^ based on normalized z-scores of averaged expressed levels per sample. For differential transcriptomic signature assessment, the SingleCellSignatureExplorer (v.3.6)^35^ was harnessed against gene sets of HALLMARK, C2, C3.TFT, and C5.BP from Molecular Signature Database (MSigDB 2023.1.Hs)^36^. Then limma (v.3.42.0) R package^37^ facilitated differential analysis of these signature scores, considering only gene sets with p.adj < 0.05 as differentially enriched transcriptome signatures (DETSs). Selected DETSs were depicted in a heatmap using pheatmap 1.0.12 package based on normalized z-scores of single-cell signature scores.

For a nuanced exploration of cancer-testis antigen (CTA) expression differences between DTCs/CTCs and their matched primary tumor cells, we examined expression metrics of CTAs listed in the CTDatabase^38^. CTAs absent in all samples were excluded from analysis. Both expression frequency (percentage of cells expressing a specific CTA) and mean expression levels of CTAs were computed. Representative CTAs, demonstrating at least a 10% elevated expression frequencies in DTCs/CTCs relative to matched primary tumor cells, were illustrated in Fig. 4g and Extended Data Fig. 8a,b through dot plots using ggcorrplot 0.1.4.1 package^39^.

### TOTT prediction using existing data

To augment the validation of the TOTT predictive accuracy across a broader spectrum of tumor types not collected in our patient cohort, we leveraged existing DTC/CTC transcriptome datasets from various prior studies. These encompassed tumor types such as liver cancer (LIHC), melanoma (SKCM), prostate cancer (PRAD), and breast cancer (BRCA)^16, 18, 40–43^. Detailed information regarding the original data sources, sample IDs, tumor types, and DTC/CTC counts can be found in Extended Data Fig. 4. For each sample, the transcriptomic data was averaged across all DTCs/CTCs, with the exception of sample DTC-B3, which utilized single-CTC transcriptome data. This consolidated data was then transformed into percentile ranks as their gene expression estimations as previously described. Subsequently, this transformed dataset served as the input for the ROM module within DTCFinder for the prediction of the tissue-of-origin and specific tumor type of the DTCs/CTCs.

### CNA inference of DTCs/CTCs via InferCNV

To juxtapose the genome-wide CNAs inferred by DTCFinder with those from established tools, we employed InferCNV to deduce CNAs from the single-cell transcriptome data of the spiked-in lung cancer cells^14^. InferCNV necessitates the designation of a cohort of non-malignant cells to serve as a reference baseline for the CNA estimation in malignant cells. To facilitate the comparison with DTCFinder’s results, we opted for the same group of immune cells used as the baseline in DTCFinder’s copy number outlier statistics. Specifically, this comprised all CD45+ immune cells, with an exclusion criterion set for those with PTRPC expression levels within the first quartile of all CD45+ cells. This baseline was utilized in InferCNV to deduce the genome-wide CNA profiles for the spiked-in cells: H1975, H1650, and A549. In the procedure, genes were arranged based on their genomic locations for each chromosome. Function infercnv::run() is called for the CNA inference of DTCs/CTCs via InferCNV with default settings except for cutoff set to 0.1 and denoise set to TRUE. A sliding window, encompassing 150 genes, was adopted to smooth the relative expression across each chromosome, thereby mitigating gene-specific expression influences. The relative expression values were subsequently centered to 1. The ceiling and floor for visualization were set using default settings, facilitating the portrayal of the data on a heatmap with a blue-red gradient, representing deletion to amplification, respectively.

### Assessment of DTCFinder’s performance on low-depth single-cell RNA sequencing data

To gauge the efficacy of DTCFinder when analyzing low-depth scRNA-seq datasets, we executed a downsampling of the original fastq files associated with DTC-N1, DTC-S1, and DTC-S2 to achieve an equivalent read depth of 5,000 reads/cell using toolkit seqtk-1.4. In detail, the original fastq file of each sample underwent downsampling into three lower-depth replicates by extracting reads randomly from the original files. The derived fastq files underwent the aforementioned preprocessing to yield feature-barcode matrices, which were subsequently used for DTC/CTC identification and TOTT tracing via DTCFinder. DTCs/CTCs discerned from the original datasets functioned as the benchmark for evaluating the sensitivity and precision of DTC/CTC identification within the downsampled datasets. The single-cell transcriptome data of DTCs/CTCs, sourced from the downsampled fastq files, was channeled into DTCFinder for TOTT predictions.

### Analysis of CNA burden across various tumor types using TCGA data

We performed a comprehensive analysis of CNA burdens in diverse tumor types, leveraging data from TCGA. The copy number segment data of primary tumor tissues, matched normal tissues and PBMCs were obtained from the Genomic Data Commons Database (https://portal.gdc.cancer.gov/). CNA burden is defined as the percentage of the tumor autosomal genome with copy number altered. To calculate CNA burden for a sample, segments manifesting copy number gains and losses were identified. The cumulative genomic length of these segments was then calculated and expressed as a percentage of the total autosomal genome size^44^.

### Statistical Analysis

Descriptive statistics, group comparisons, and correlation tests were performed using GraphPad PRISM 10 (GraphPad Software, Inc) unless noted elsewhere.

## Acknowledgements

We thank the following agencies and foundations for support: NIH grant R33 CA256112 (to W.W.); Andy Hill CARE Fund (to W.W.); National Natural Science Foundation of China (No. 22374027, to Q.S.) and Program of Shanghai Academic/Technology Research Leader (No. 23XD1403900, to Q.S.). Q.S. was also sponsored by BeiGene.

## Author contributions

W.W. and Q.S. conceived and supervised the study. X.Y., Z.W., Z.L. and Y.Z. performed the research. X.Y. and W.W. developed the algorithm. X.Y., Z.W. W.W. and Q.S. analyzed the data. Z.L. and S.L. provided resources for this research. W.W., X.Y. and Q.S. wrote the paper with input from other authors. All authors read and approved the final paper.

---

**Extended Data Fig. 1.**
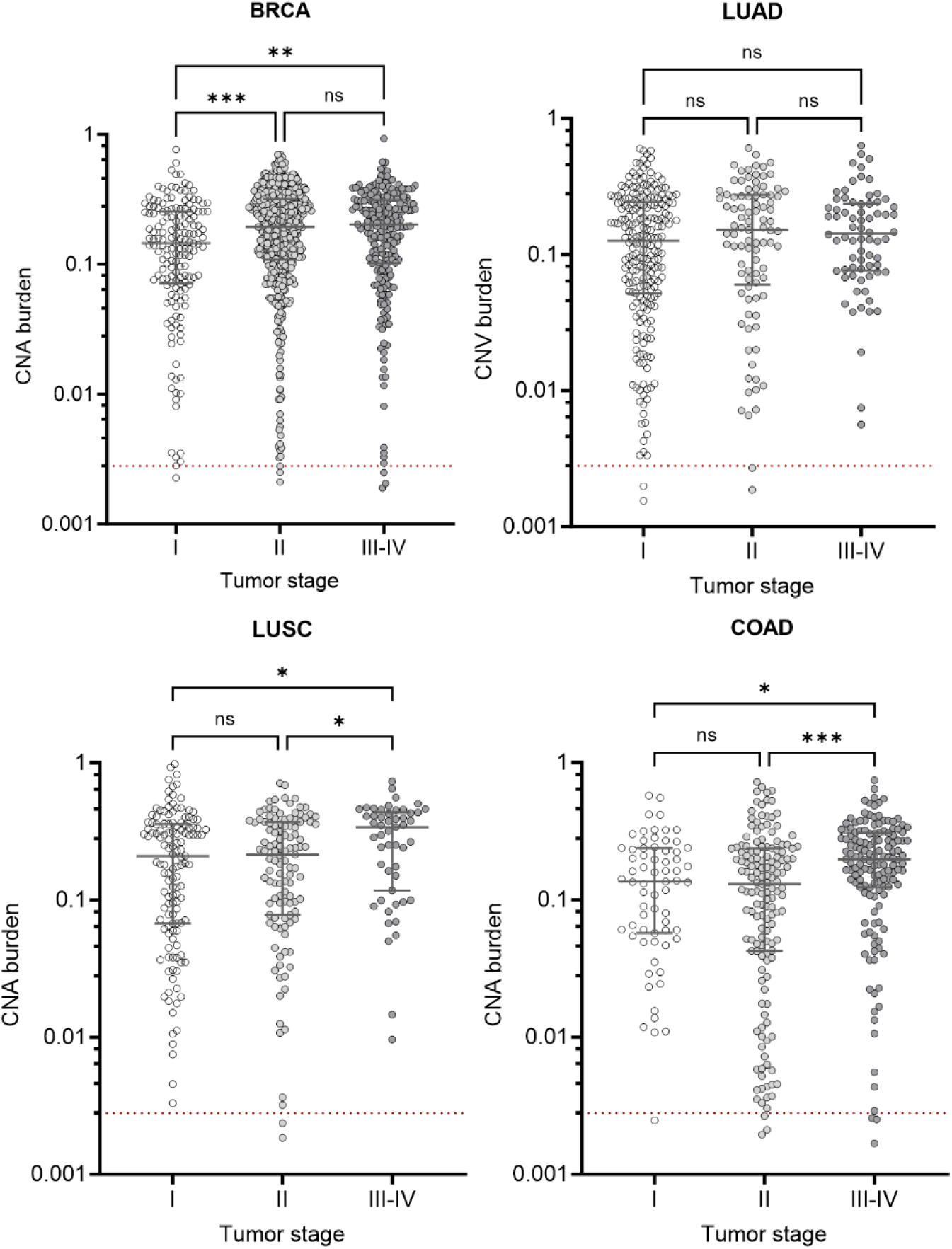
Evaluation of CNA burdens across varied stages for four exemplary tumor types. The depicted CNA burdens for distinct tumor samples from TCGA, classified according to their respective stages, are subjected to pairwise comparisons employing the Kruskal-Wallis test, supplemented by Dunn’s test for corrections pertaining to multiple comparisons. The red dash line indicates the average CNA burden of normal immune cells (*p<0.05, **p<0.005, ***p<0.0005, ns: not significant).

**Extended Data Fig. 2.**
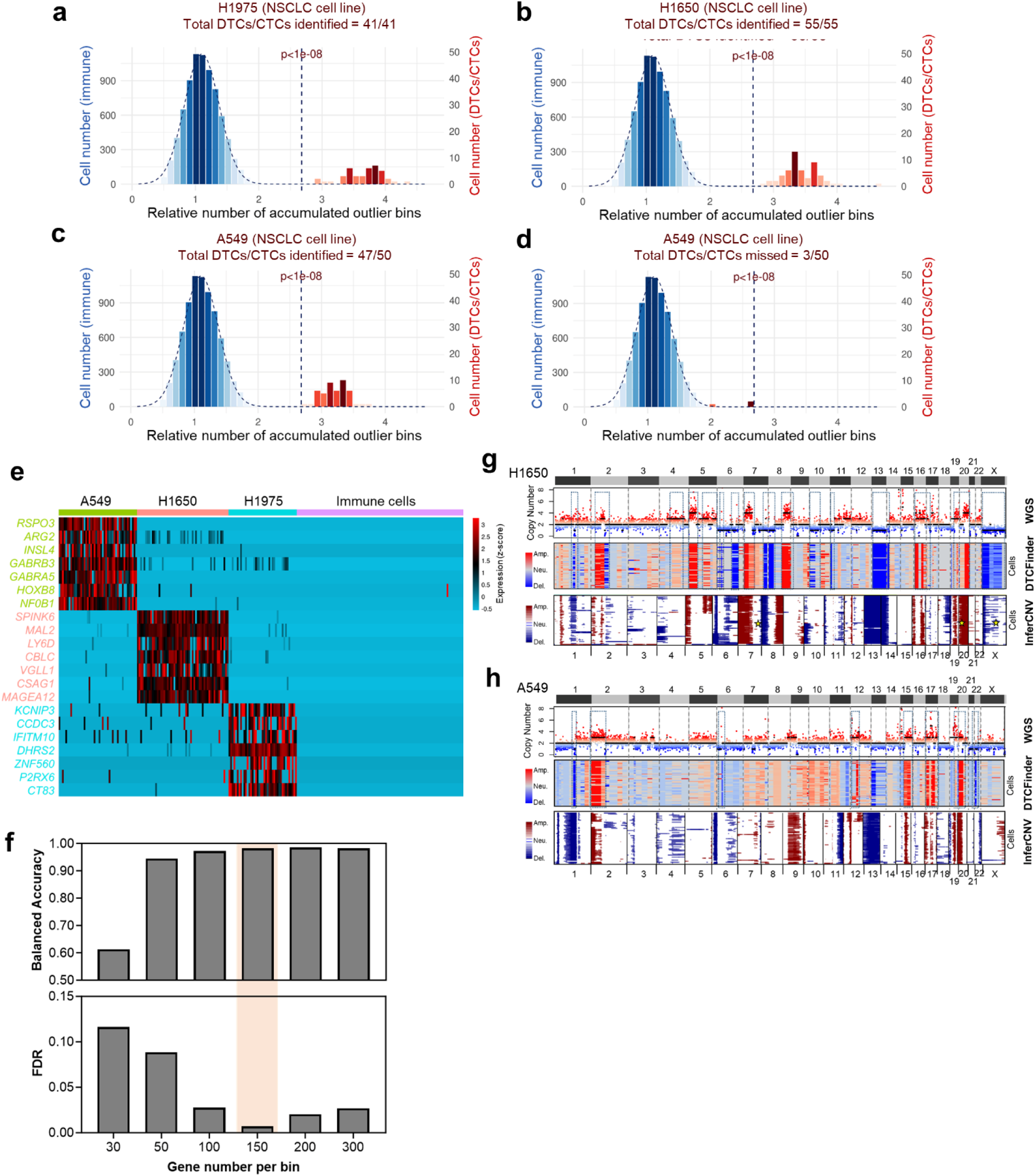
Assessment of DTCFinder performance by the cell line spike-in sample. **a-c,** Histograms of accumulated outlier bins (AOBs) for baseline immune cells, depicted with Gaussian fittings, juxtaposed with spike-in H1975 (**a**), H1650 (**b**), and A549 (**c**) cells as identified by DTCFinder. Dashed lines represent the AOB cut-off p-value of 10^-8^. Ratios of identified to actual present tumor cells are displayed above each histogram for individual cell lines. **d.** Histogram illustrating AOBs for baseline immune cells (blue) and spike-in A549 cells (red) not detected by DTCFinder, highlighting proximity of AOBs of missed A549 cells to the DTC/CTC identification cut-off threshold. **e.** Normalized z-scores represent expression levels of cell line-specific markers across spike-in A549, H1650, H1975 cells, and randomly selected immune cells, with color coding for cell line-specific gene markers: green for A549, red for H1650, and blue for H1975. **f.** DTCFinder’s performance at varying bin sizes is illustrated by the balanced accuracy and false discovery rate (FDR) of DTC identification, establishing 150 genes/bin as the default setting due to near-perfect balanced accuracy and minimal FDR. **g, h.** Comparative illustrations of genome-wide CNAs as measured by LC-WGS and inferred by DTCFinder and InferCNV for H1650 (**g**) and A549 (**h**). Genomic segments manifesting consistent CNAs are encircled by blue dashed boxes. Copy number profiles at specific genomic loci, accurately inferred by DTCFinder but imprecisely inferred by InferCNV, are marked by yellow stars.

**Extended Date Fig. 3.**
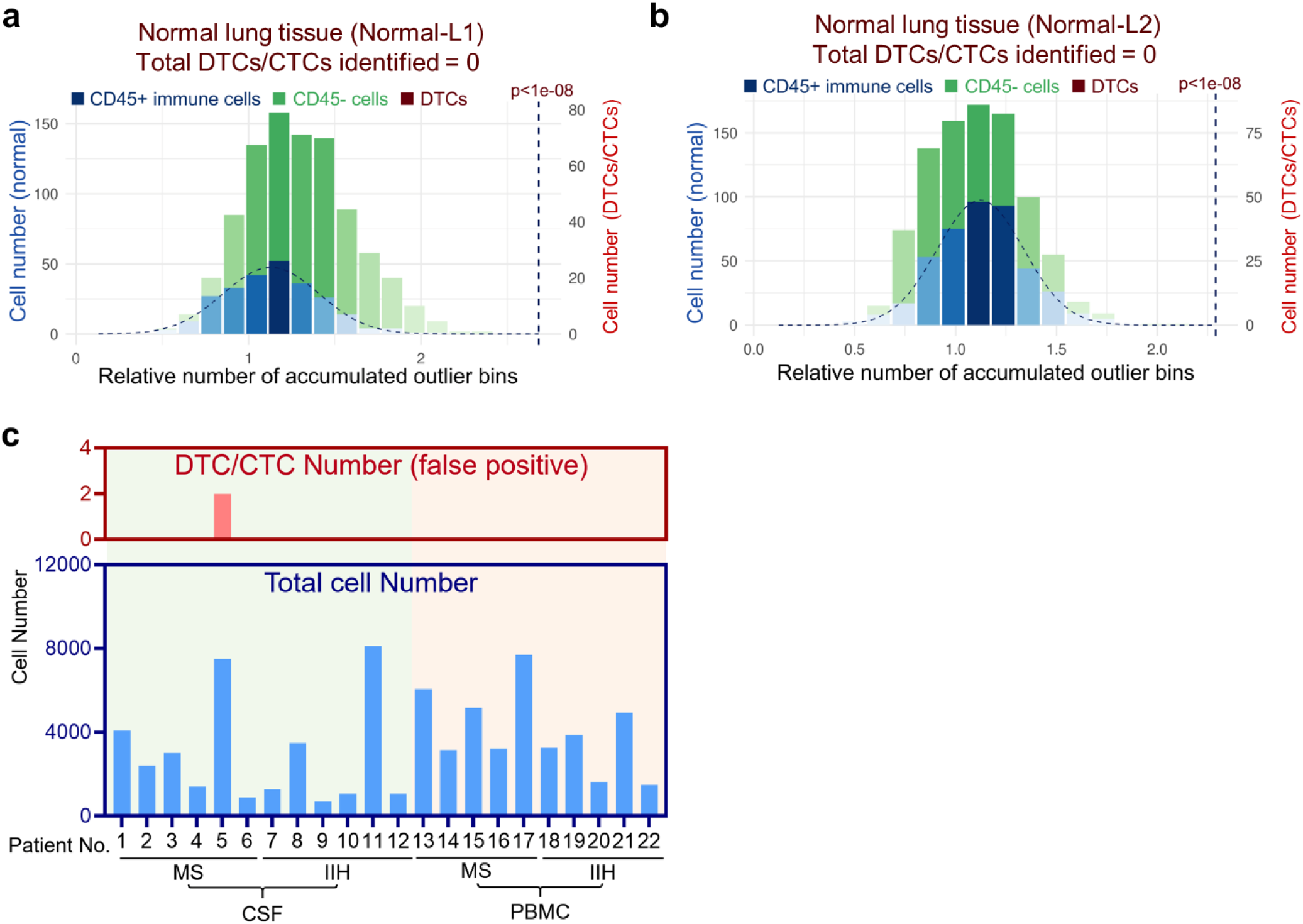
Examination of DTCFinder’s specificity with normal lung tissues and non-cancerous liquid biopsy samples. **a, b.** Histograms depict accumulated outlier bins (AOBs) for baseline immune cells (blue) with Gaussian fitting contrasted with non-immune epithelial cells from normal lung tissues (green) from patients Normal-L1 (**a**) and Normal-L2 (**b**). No cells in these samples surpassed the threshold to be falsely identified as DTCs/CTCs. **c.** Total cell number and false positive cells identified by DTCFinder in PBMC and CSF samples collected from a cohort of patients diagnosed with multiple sclerosis (MS) and idiopathic intracranial hypertension (IIH).

**Extended Date Fig. 4.**
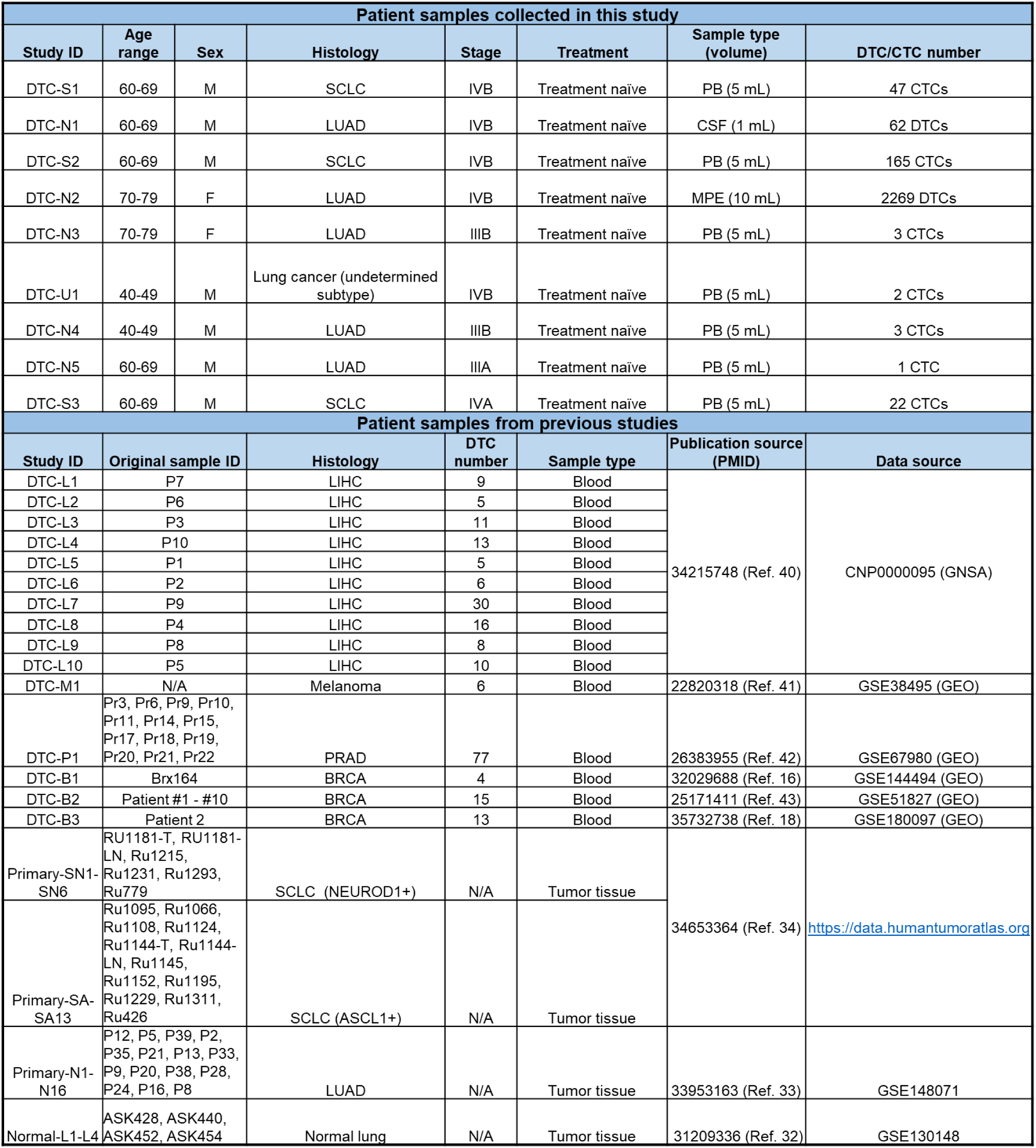
Summary of patient samples analyzed in this study. Every sample is assigned a unique study ID. The clinical metadata, sample type, and DTC/CTC number identified are itemized for each patient sample gathered in this study. For the analysis of samples from previous studies, the original sample IDs, pertinent sample information, and data sources are cataloged for each patient sample. PB: peripheral blood, CSF: cerebrospinal fluid, MPE: malignant pleural effusion. Key for histology: LUAD, lung adenocarcinoma; SCLC, small cell lung cancer; LIHC, liver hepatocellular carcinoma; PRAD, prostate adenocarcinoma; BRCA, breast carcinoma.

**Extended Date Fig. 5.**
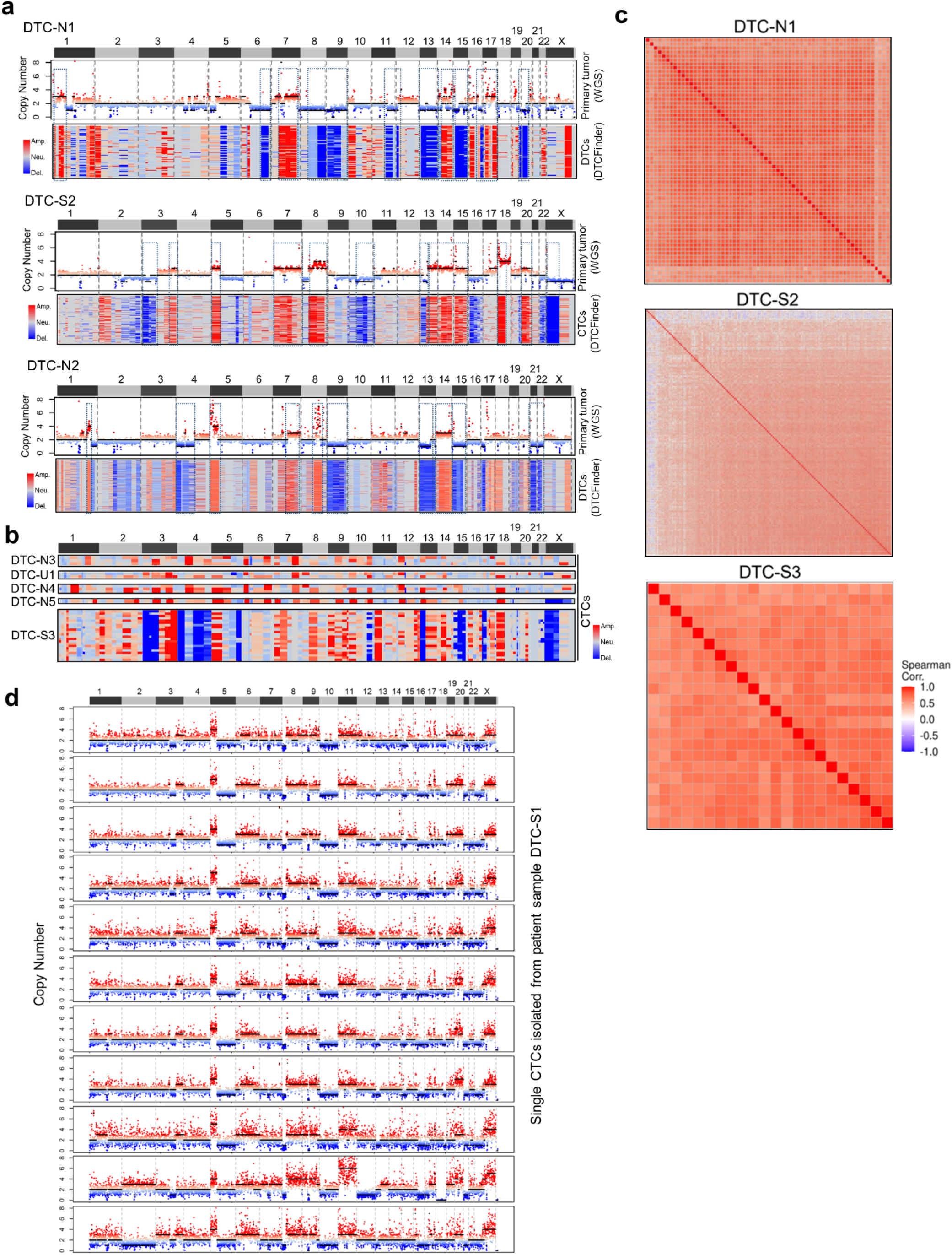
In-depth analysis of genome-wide CNA by DTCFinder for DTCs/CTCs identified in liquid biopsy samples from lung cancer patients in this study. **a.** Alignments of genome-wide CNAs of DTCs/CTCs inferred by DTCFinder with those of paired primary tumor tissues ascertained by LC-WGS are illustrated for patients DTC-N1, DTC-S2, and DTC-N2. Genomic segments with consistent CNAs are outlined by blue dashed boxes. **b.** Depictions of inferred genome-wide CNA profiles by DTCFinder for DTCs/CTCs from patients DTC-N3, DTC-U1, DTC-N4, DTC-N5, and DTC-S3. **c.** Heatmaps showing pairwise Spearman’s correlations of inferred CNAs across all identified DTCs/CTCs in patients DTC-S1, DTC-S2, and DTC-S3, respectively. **d.** Genome-wide CNAs measured by LC-WGS for 11 randomly selected single CTCs isolated from the peripheral blood of patient DTC-S1 reveals highly consistent (correlated) CNA patterns across different CTCs, corroborating the high correlations of the inferred CNA profiles.

**Extended Date Fig. 6.**
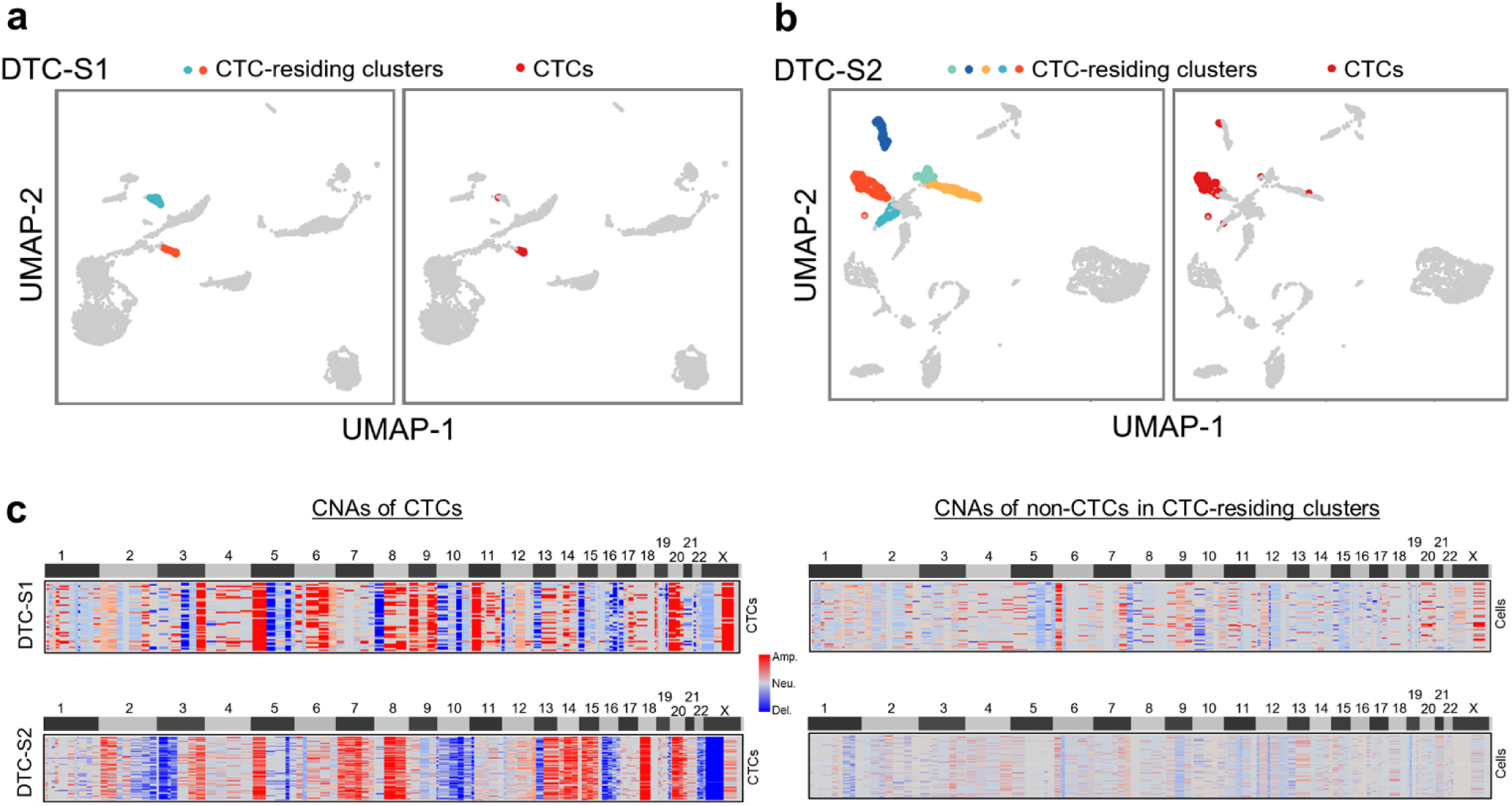
Evaluation of CTC identification by direct clustering analysis. **a, b.** The Louvain clustering method applied to scRNA-seq data from patients DTC-S1 (**a**) and DTC-S2 (**b**) is illustrated, with CTC-residing clusters (left) and CTCs (right) color-coded and represented in UMAP projections. **c.** The inferred genome-wide CNAs by DTCFinder for CTCs and remaining cells within the CTC-residing clusters are displayed for patients DTC-S1 (top) and DTC-S2 (bottom).

**Extended Date Fig. 7.**
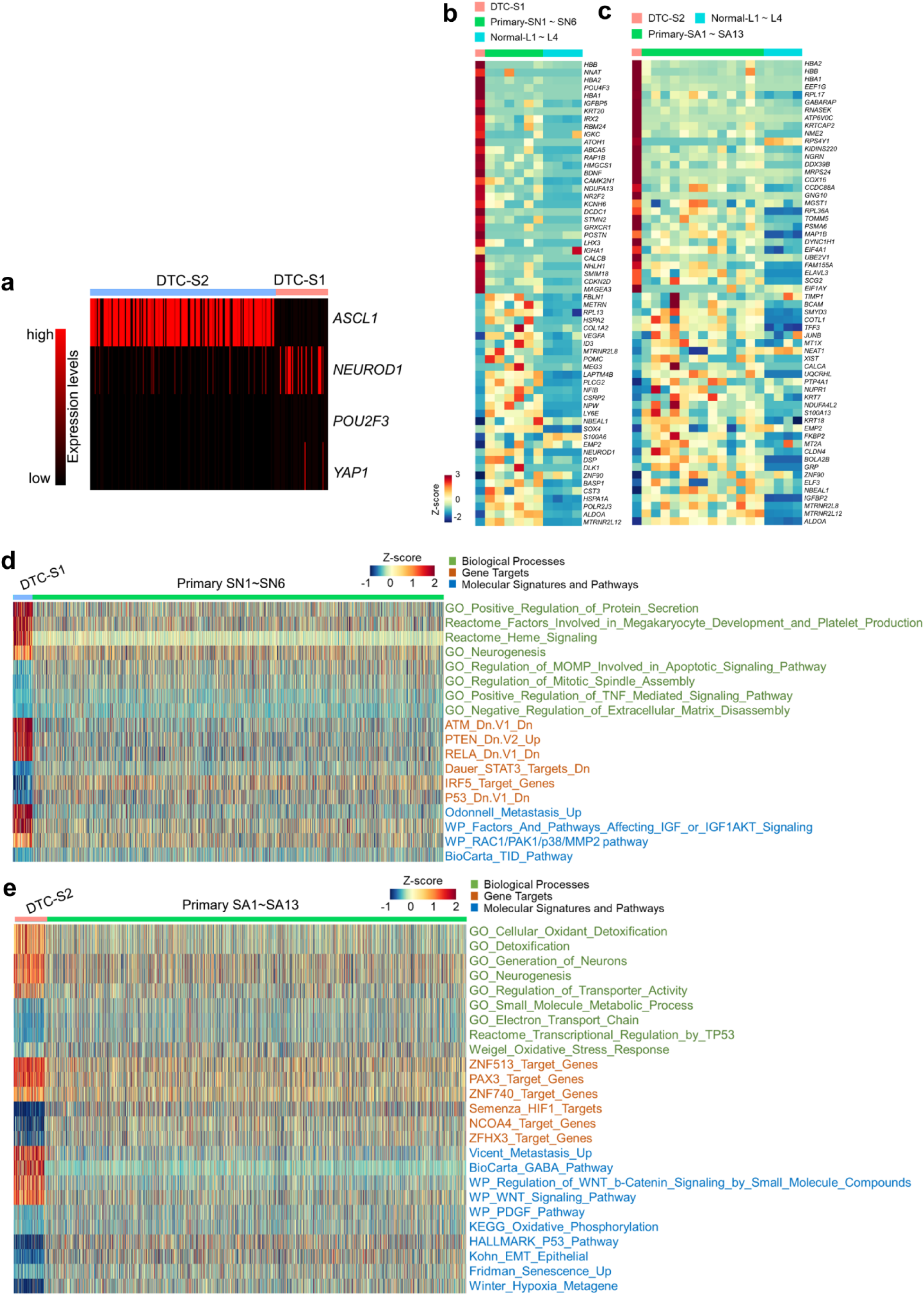
Transcriptome characterization of DTCs/CTCs and corresponding primary tumors. **a.** A heatmap illustrates the expression levels of marker genes delineating different SCLC subtypes across identified CTCs from peripheral blood samples of patients DTC-S1 and DTC-S2. **b, c.** The top and bottom 30 differentially expressed genes in CTCs from SCLC patients DTC-S1 (**b**) or DTC-S2 (**c**) are juxtaposed with primary tumor cells from patients harboring the same SCLC subtype as well as normal lung samples, with the averaged gene expression levels of each sample represented based on normalized z-scores. **d, e.** Representative differentially enriched transcriptome signatures between CTCs from SCLC patients DTC-S1 (**d**) or DTC-S2 (**e**) and primary tumor cells from patients with the same SCLC subtype are color-coded and illustrated with normalized z-scores. The data bars for CTCs are magnified fivefold for enhanced clarity.

**Extended Date Fig. 8.**
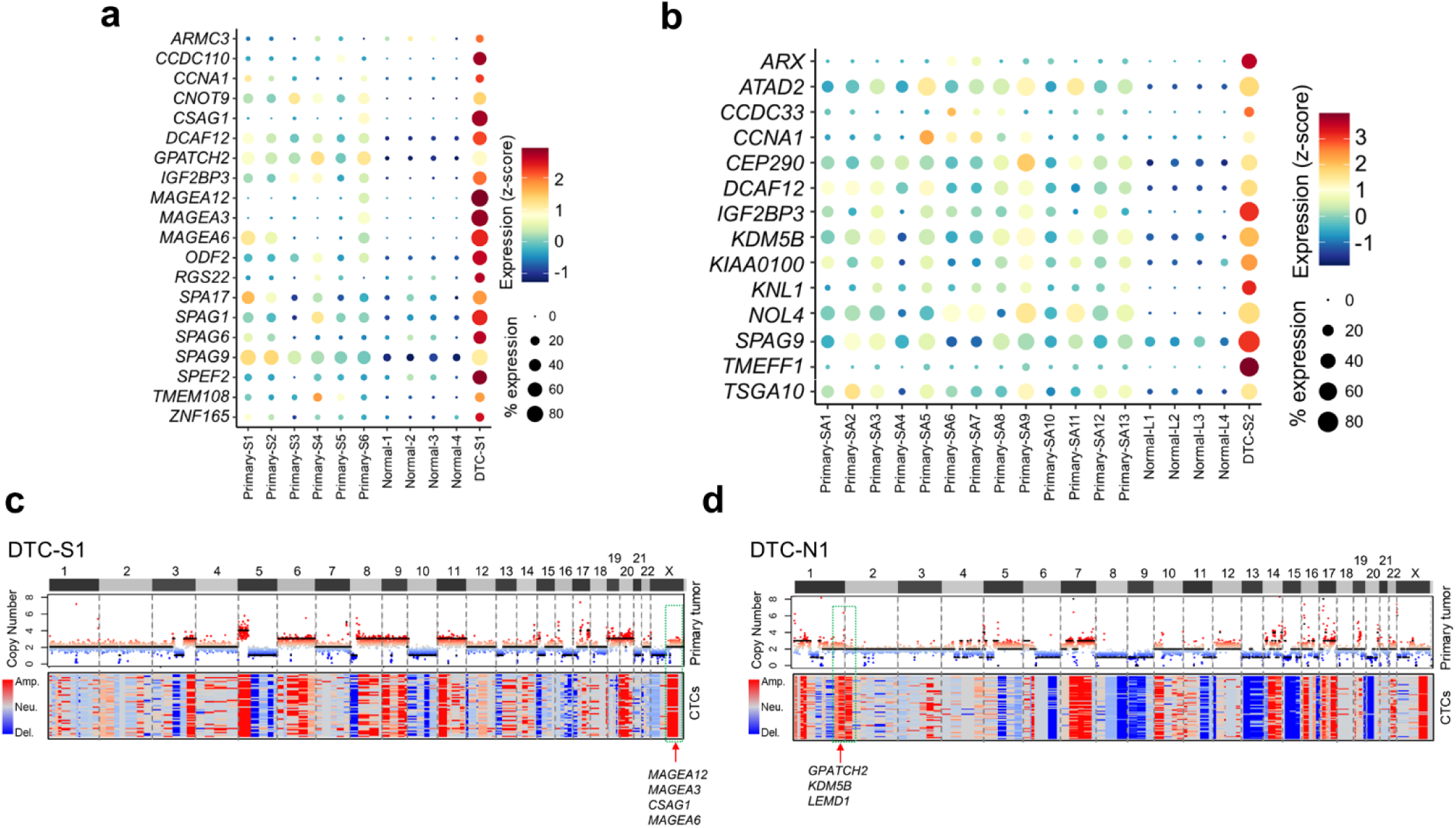
Increased expression of cancer-testis antigens (CTAs) in DTCs/CTCs. **a.** Comparative analysis of expression levels and frequencies of representative CTAs amongst CTCs from SCLC patient DTC-S1, primary tumor cells from six patients of the same SCLC subtype, and epithelial cells from four normal lung tissues. **b.** Comparative analysis of expression levels and frequencies of representative CTAs between CTCs from SCLC patient DTC-S2, primary tumor cells from 13 patients with the same SCLC subtype, and epithelial cells from 4 normal lung tissues. For both (**a**) and (**b**), normalized z-scores show averaged CTA expression across single cells within each sample, with circle sizes indicating expression frequencies (i.e. the percentage of cells expressing the CTA). **c, d.** Genomic segments with DTC/CTC-specific CNAs coinciding with loci of overexpressed CTAs are outlined by green dashed boxes, with associated CTAs enumerated below for patients DTC-S1 (**c**) and DTC-N1 (**d**)

**Extended Date Fig. 9.**
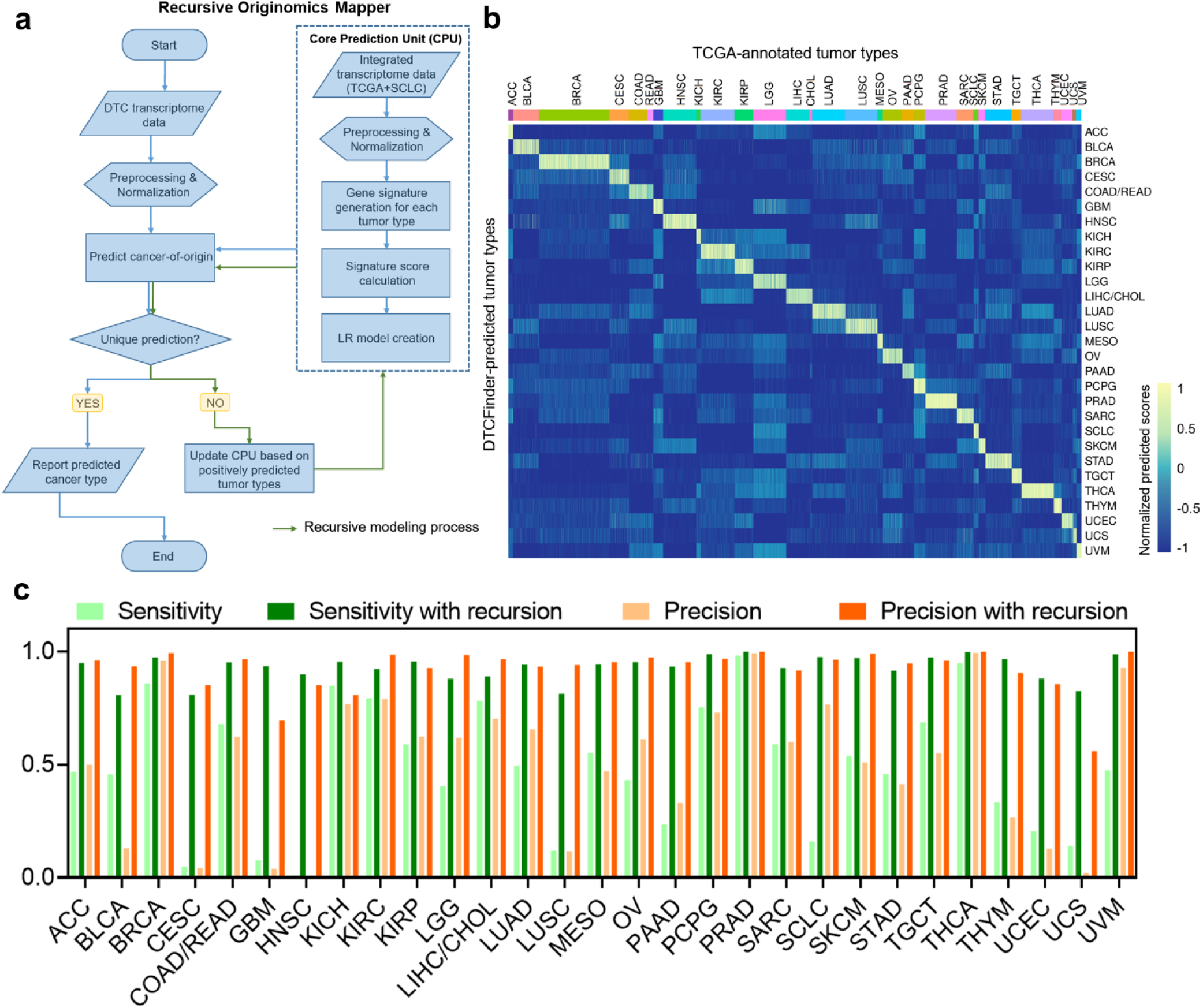
Design and evaluation of DTCFinder’s model for tissue-of-origin and tumor type (TOTT) tracing. **a.** A flowchart elucidates the construction of the recursive logistic regression (LR)-based Originomics Mapper, designed to trace TOTT utilizing single-cell transcriptomic data of DTCs/CTCs. **b.** The representation of TOTT prediction outcomes, utilizing transcriptome data from patients across 31 solid tumor types within the TCGA database, is conducted in a 10-fold cross-validation setting. Scores exceeding zero indicate a positive prediction corresponding to a specific tumor type. **c.** Sensitivities and precisions of TOTT predictions rendered by DTCFinder spanning 31 TCGA solid tumor types with or without the integration of the recursive process.

**Extended Date Fig. 10.**
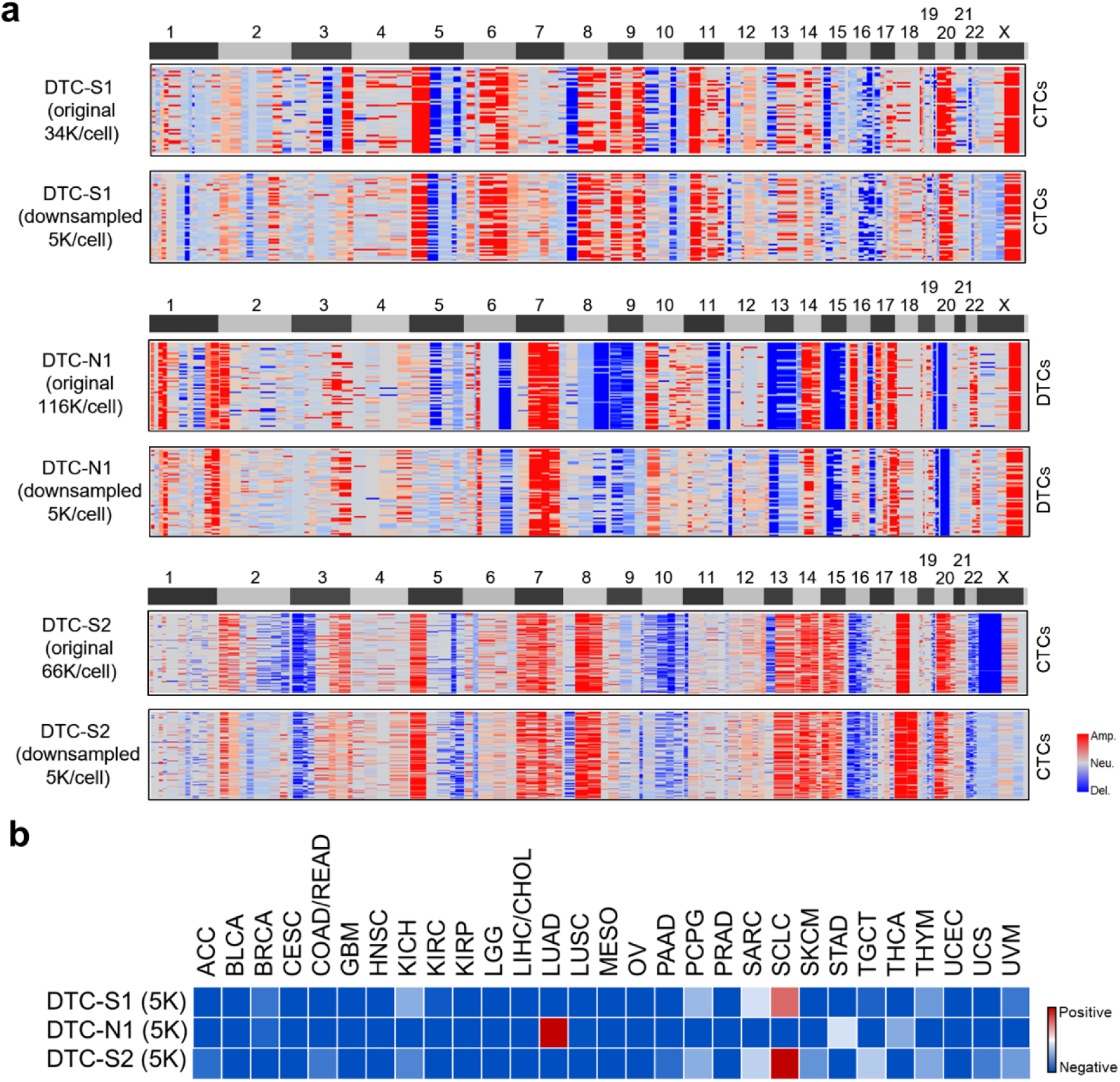
Performance of DTCFinder on low-depth sequencing data. **a.** Comparison of inferred genome-wide CNA profiles between original sequencing depth and low-depth (5K reads/cell) data across three lung cancer samples. **b.** DTCFinder’s TOTT prediction using downsampled, low-depth (5K reads/cell) data across three lung cancer samples.

